# Insulin-Induced Conformational Changes in the Full-Length Insulin Receptor: Structural Insights Gained from Molecular Modeling Analyses

**DOI:** 10.1101/2020.09.01.278812

**Authors:** Yong Xiao Yang, Peng Li, Pan Wang, Bao Ting Zhu

## Abstract

Insulin receptor plays an important role in regulation of energy metabolism. Dysfunction of insulin receptor (IR) can lead to many disease states, such as diabetes mellitus. Deciphering the complex dynamic structure of human IR and its mechanism of activation would greatly aid in understanding IR-mediated signaling pathways and in particular, in designing new drugs (including nonpeptidal insulin analogs) to treat diabetes mellitus. Experimental evidence about IR structure has been gradually obtained by biologists over the past three decades. Based on the available experimental structures of IR in different states, here we employ molecular modeling approach to construct the full-length IR structures in different states and model its structural and conformational changes during insulin-induced IR activation. Several key possible intermediate states are constructed based on structural alignment, rotation and computational modeling. Based on the structures of the full-length IR in different states, it appears that there are two possible conformational transition pathways: one is symmetric, and the other one is asymmetric. Structural changes and motions of different domains of the full-length IR along the pathways are analyzed. The role of insulin binding to IR in facilitating the conformational transition of the receptor is modeled. Information and insights derived from our present structural modeling analyses may aid in understanding the complex dynamic, structural and conformational changes during the process of IR activation.

## INTRODUCTION

Insulin receptor (IR), a member of the receptor tyrosine kinase family, is of great importance in regulating many important cellular functions, including metabolism (in particular glucose metabolism and homeostasis), cellular growth, differentiation and survival [1–3]. Dysfunction of the IR and its associated signaling pathways contributes critically to the pathogenesis of many disease states, such as type 2 diabetes mellitus and Alzheimer’s disease [4]. Understanding the complex structure of IR and its activation is believed to be of fundamental importance in biology and medicine, and will aid significantly in the rational design of specific drugs for related disease conditions.

IR was sequenced by Ebina *et al.* in 1985 [5]. It is a native α2β2 tetramer [6], and contains an extracellular ectodomain, a transmembrane domain, and an intracellular tyrosine kinase domain. Since the determination of the first structure of tyrosine kinase domain by Hubbard *et al.* in 1994 [7], experimental structural biologists have resolved the structures of different segments of human IR [8, 9]. Especially, the structures of the ectodomain in different states have been reported in the past 15 years [9], which offers important insights into the molecular mechanism and structural basis of IR activation. Many research groups have been working on the structures of IR’s ectodomain, as it is directly involved in the binding interaction with its endogenous ligand insulin. Ward and Lawrence *et al*. have resolved a number of crystal and cryo-EM structures of IR’s ectodomain in the past 15 years [10–15]. In recent years, research teams led by Scapin, Gutmann and Bai have also resolved several cryo-EM structures of IR’s ectodomain in different states [16–18]. Scapin *et al.* and Gutmann *et al.* proposed two similar models of IR activation which mostly represent a symmetric-asymmetric-symmetric conformational transition pathway [16, 17], although some differences are noted between the first suppositional symmetric states in these two models. These two earlier studies were partly based on the structure of one chain of the ectodomain reported by Croll *et al*. [12] (PDB code: 4ZXB). Bai *et al.* also proposed a working model of insulin-induced IR activation which is a symmetric conformational transition pathway [18]. Additionally, Li *et al.* in 2014 [19] determined the structure of the transmembrane domain by using NMR, and in the ensuing year, Cabail *et al.* reported the tyrosine kinase domain as a functional dimer at the active states [20]. In all these models, the initial states were speculated, and there appears to be no complete experimental structure that represents the corresponding conformations. Also, there lacks detailed analysis of the conformational changes of full-length IR during the process of insulin-induced IR activation.

In the present work, efforts are made to model and assemble the different states and structures of the full-length IR based on the experimental data or structural information available in the literature. In addition, attempts are made to model and analyze the conformational changes during the process of insulin-induced activation of the full-length IR.

## RESULTS AND DISCUSSION

### Structures of the full-length IR in different states

IR contains two chains with the same amino acid sequence and domain arrangements, and combination of domains from two chains jointly form some of the structures, such as the insulin-binding sites. In this work, when IR’s one chain is mentioned, the other chain is regarded as the partner chain (or simply as “the partner”); and if some of the domains of one chain are labeled as L1, αCT and Fnш-1, their counterpart domains of the other chain are labeled as L1’, αCT’ and Fnш-1’, respectively.

In an earlier study by Gutmann *et al.* [21], there are mainly three conformations of the full-length IR, namely, U-shaped, II-shaped (asymmetric), and T-shaped, which were observed by single particle electron microscopy using glycosylated full-length human IR reconstituted into lipid nanodiscs. It is known that the U-shaped conformation exists in the absence of insulin (**Fig. 1A**), and the II-shaped and T-shaped conformations exist in the presence of insulin (**Fig. 1C** and **1D**) [21]. Additionally, the full-length IR in two lipid nanodiscs also presents a T-shaped conformation in the presence of insulin, which likely represents an intermediate state (**Fig. 1B**) [21]. Different groups of cryo-EM images obtained by Gutmann *et al.* [21] are shown in **Fig. S1**, which are included for reference and comparison, whereas the four structures in **Fig. 1** are used as main references for constructing the structures of full-length IR based on some of the available experimental structural information in the literature.

**Figure 1.**
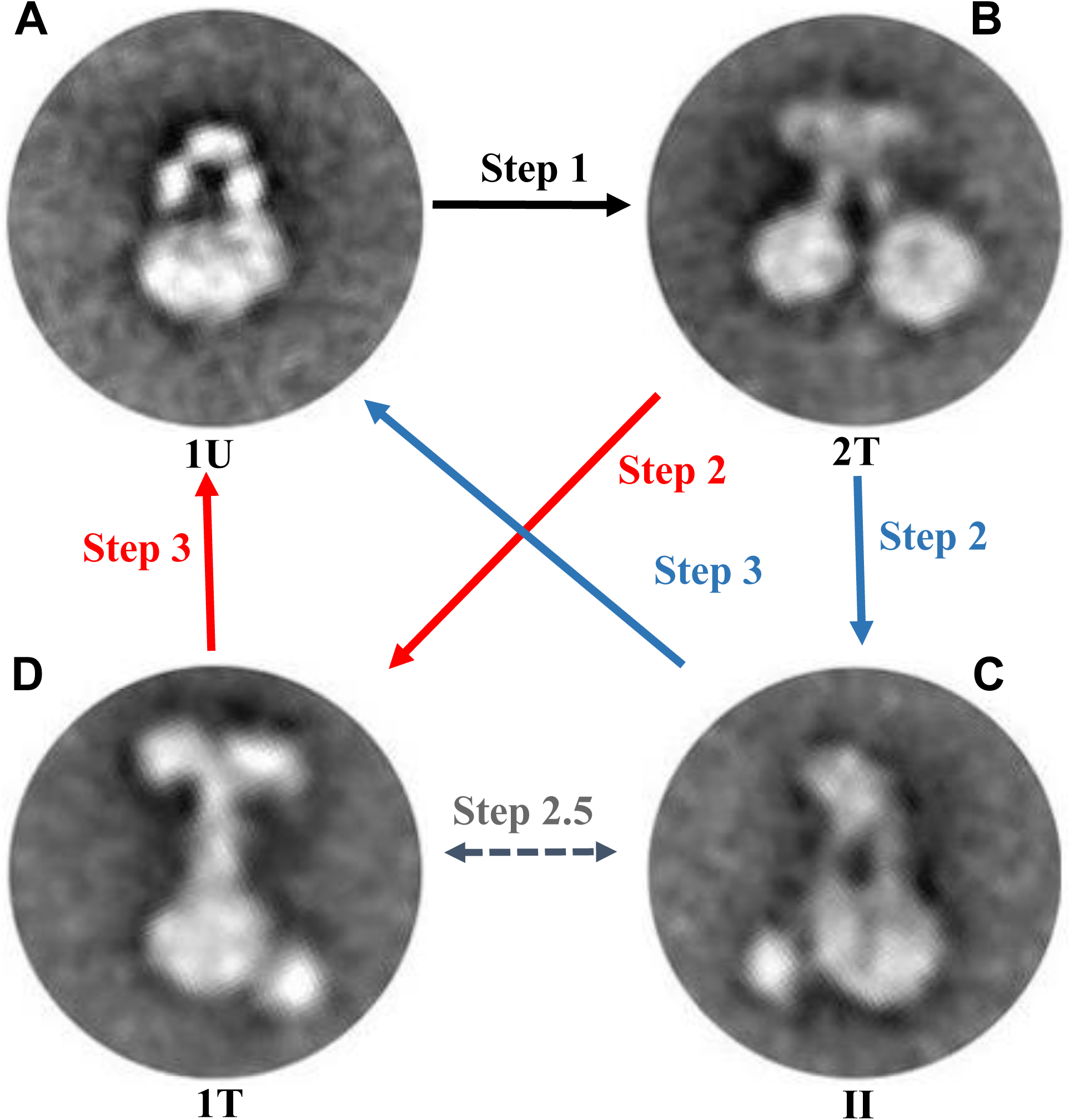
Cryo-EM images of the conformations of the full-length IR observed by Gutmann *et al.* [21]. **A.** In the absence of insulin, the full-length IR presents an (inverted) U-shaped conformation in single lipid nanodics, which is considered to be the apo form (auto-inhibited state). **B.** In the presence of insulin, the full-length IR presents a T-shaped conformation in two lipid nanodiscs, which may be a symmetric intermediate state. **C.** In the presence of insulin, the full-length IR presents an II-shaped conformation in single lipid nanodisc, which may be an asymmetric active state. **D.** In the presence of insulin, the full-length IR presents a T-shaped conformation in single lipid nanodisc, which may be a symmetric active state.

There are two types of insulin binding sites in IR structures [17, 18], *i.e.*, type 1 binding site (T1BS) and type 2 binding site (T2BS). The complete T1BS involves the L1 domain of one chain and the αCT’ and Fnш-1’ domains of the partner chain. T2BS is located on the side of the Fnш-1 domain. Different domains of IR are shown in **Fig. S2**: the full-length IR (one chain) (**Fig. S2A**), the signal peptide (**Fig. S2B**), L1 (leucine-rich repeat domain 1) (**Fig. S2C**), CR (cysteine-rich region) (**Fig. S2D**), L2 (leucine-rich repeat domain 2) (**Fig. S2E**), Fnш-1 (fibronectin type-ш domain 1) (**Fig. S2F**), Fnш-2a (fibronectin type-ш domain 2a) (**Fig. S2G**), ID-α (insert domain of the α chain) (**Fig. S2H**), αCT (C-terminal domain of the α chain) (**Fig. S2I**), ID-β (insert domain of the β chain) (**Fig. S2J**), Fnш-2b (fibronectin type-ш domain 2b) (**Fig. S2K**), Fnш-3 (fibronectin type-ш domain 3) (**Fig. S2L**), TM and JM (transmembrane domain and juxtamembrane domain, respectively) (**Fig. S2M**), TK (tyrosine kinase) (**Fig. S2N**), and βCT (C-terminal domain of the β chain) (**Fig. S2O**).

#### Construction of the apo form of IR

The U-shaped conformation of IR is also known as an auto-inhibited state (*i.e.*, the apo form) [17, 18, 21]. In 2006, Ward *et al.* determined the crystal structure of the first three domains (L1∼CR∼L2 domains) of IR which forms a L1∼CR∼L2∼L1’∼CR’∼L2’ homodimer [10] (PDB code: 2HR7, left panel of **Fig. S3A**). Almost the same time, Ward and Lawrence *et al.* reported a crystal structure of the IR’s ectodomain [11] with an improved resolution at 3.30 Å [12] (PDB code: 4ZXB, right panel of **Fig. S3A**). The L1∼CR∼L2 domains are a shared part in the two incomplete experimental structures of IR determined by Ward and his colleagues in 2006 [10–12] (**Fig. S3A**).

To obtain the complete U-shaped ectodomain based on the L1∼CR∼L2∼L1’∼CR’∼L2’ homodimer conformation [10] (left panel of **Fig. S3A**), the shared L1∼CR∼L2/L1’∼CR’∼L2’ domains are superimposed with VMD [22] and the missing Fnш and αCT domains are placed into the corresponding position of the homodimer structure. Because the angle between the L1∼CR∼L2 domains and the Fnш and αCT domains [12] (right panel of **Fig. S3A**) does not permit the formation of the U-shaped conformation of the ectodomain, the Fnш and αCT domains are thus rotate with VMD [22] to properly adjust the angle such that the U-shaped ectodomain can be formed. Then, the transmembrane domain, juxtamembrane domain and tyrosine kinase domain are added to the U-shaped ectomdomains. The missing segments such as ID-α and ID-β in the ectodomain are constructed by SWISS-MODEL [23]. Because no templates are available, the signal peptide (27 amino acids) and part of the βCT domains (73 amino acids) are also predicted using Confold2 [24]. The secondary structures and residue-residue contacts are predicted using SPOT-1D [25] and SPOT-Contact [26], respectively. Then all experimental and predicted structures are assembled into a complete full-length IR (**Fig. 2A**). The main characteristics of this full-length structure in its U-shaped conformation are that the L1 domain makes contacts with the L1’ and L2’ domains of the partner, but not with the Fnш-1’ domain of the partner. Also, the two-T2BSs for insulin on the two Fnш-1 domains are exposed to the solvent and appear to be ready for insulin binding. It is of note that in this structure model, the distance between the two Fnш-3 domains is quite long such that the two transmembrane-juxtamembrane-tyrosine kinase domains are separated.

**Figure 2.**
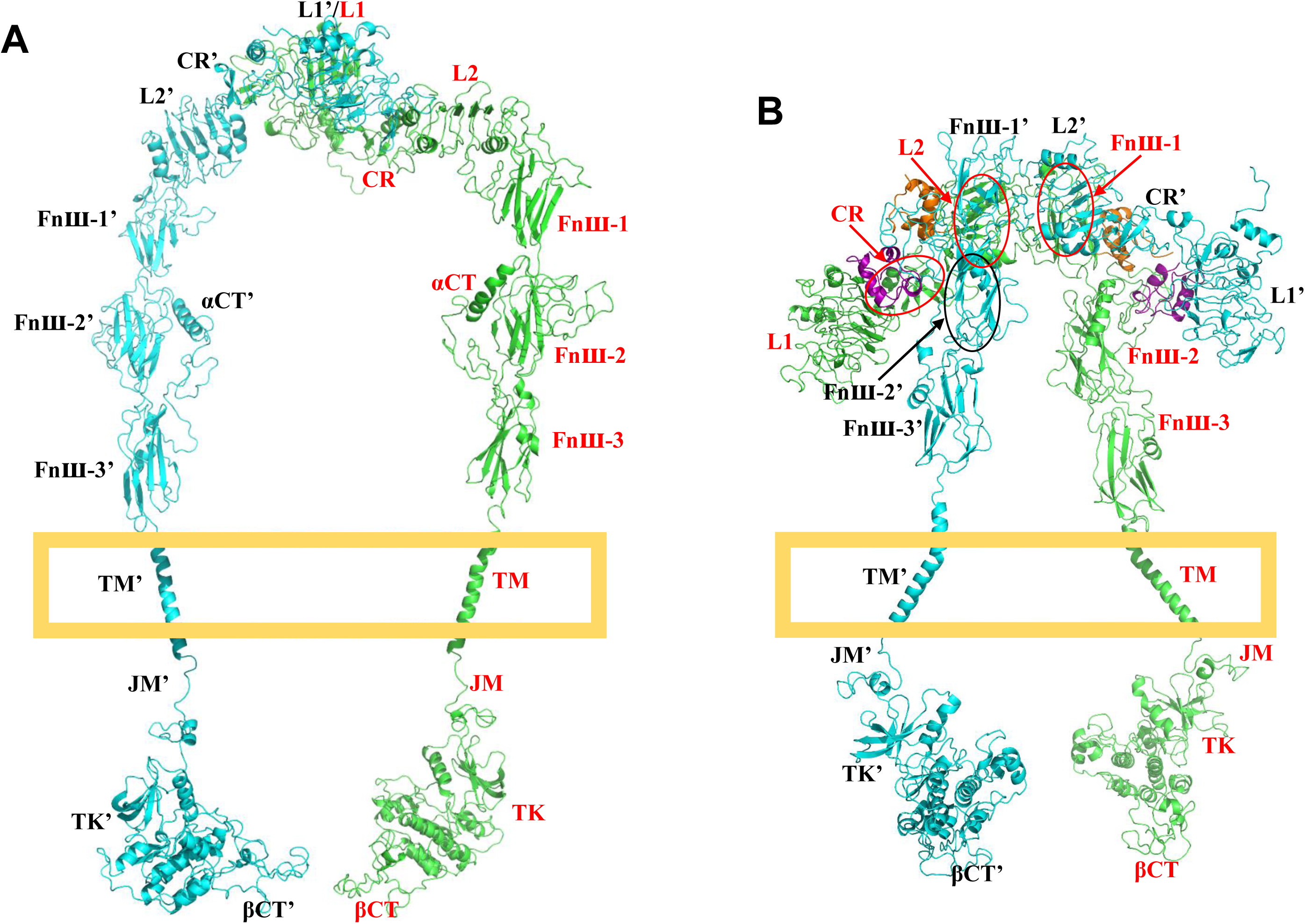
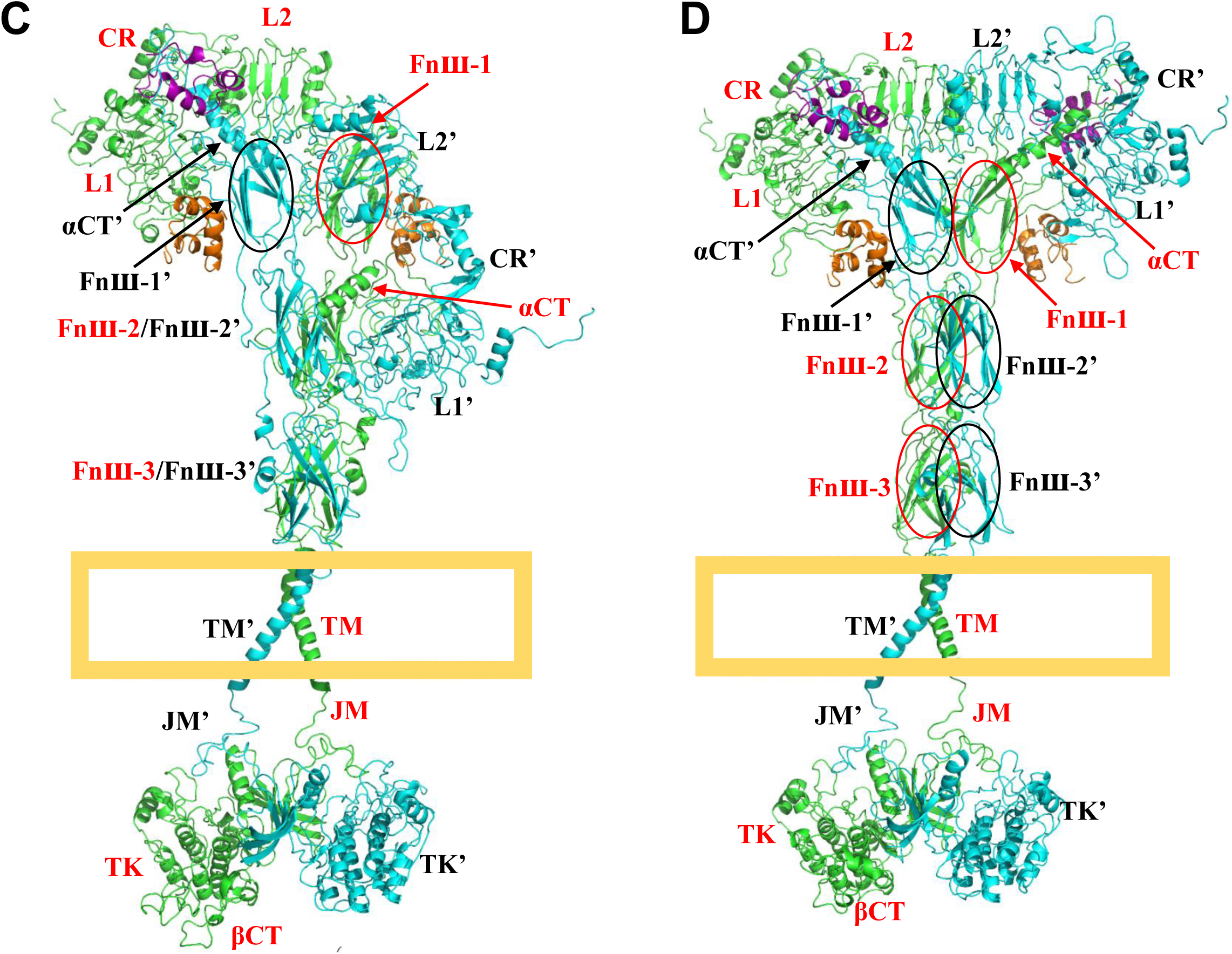
Structures of the constructed full-length IR at four representative states. **A.** The Apo form (auto-inhibited state; based on PDB code: 2HR7, 4ZXB, 2MFR, 4XLV). **B.** Symmetric intermediate state (based on PDB code: 5KQV, 6HN4, 4ZXB, 2MFR, 4XLV). **C.** Asymmetric active state (based on PDB code: 6HN4, 6HN5, 4ZXB, 2MFR, 4XLV). **D.** Symmetric active state (based on PDB code: 6PXV, 6SOF, 2MFR, 4XLV). The insulins at the incomplete or complete T1BSs are colored in purple, and insulins at the T2BSs colored in orange. Each pair of arrow and ellipse in red points to a domain in one partner chain which is colored in green; likewise, the pair of arrow and ellipse in black points to a domain in the other partner chain which is colored in cyan.

We notice that in some of the IR activation models reported in earlier studies [9, 17, 18], the initial state or the auto-inhibited state does not share a similar U-shaped conformation as indicated by our reconstructed model. In our constructed initial state of the full-length IR (shown in **Fig. 2A**), it appears that there is no direct contact between the Fnш-1 domain and the L2’ domain of the partner; but in the earlier proposed IR-activation models [9, 17, 18], the Fnш-1 domain in the initial state makes contacts with the L2’ domain of the partner, forming a different U-shaped conformation.

It is of note that there are two experimental structures presently available in the literature which report that the Fnш-1 and Fnш-1’ domains make contacts with the L2’ and L2 domains of the partners, respectively. The first one was initially determined by Ward and Lawrence *et al.* in 2006 [11], but the structure deposited in Protein Data Bank [27] was later substituted with another structure with only one chain of IR [12] (PDB code: 4ZXB; right panel of **Fig. S3A**). This information has served as reference for the initial state in the IR activation models proposed later by other researchers [9, 17, 18]. The second one was determined by Ward, Lawrence and their colleagues in 2013 [13], which is a crystal structure of the L1∼CR∼L2∼Fnш-1∼L1’∼CR’∼L2’∼Fnш-1’ dimer revealing two insulins bound inside the two incomplete T1BSs (PDB code: 5KQV; left panel of **Fig. S3B**). This important structural information is also used in this study to construct the intermediate state of IR (detailed information is provided below).

#### Construction of the intermediate state of IR

In 2014, Ward, Lawrence and colleagues further reported a crystal structure of the insulin∼L1∼CR complex with a higher resolution [14] (PDB code: 4OGA; right panel of **Fig. S3B**). The structure (shown in **Fig. S3B**) likely represents parts of the proposed intermediate state in the presence of insulin (described below), which is a T-shaped conformation (in two lipid nanodiscs) (**Fig. 1B**) as reported by Gutmann *et al.* [21].

To assemble the full-length IR in its possible intermediate state(s), the missing domains and segments are added according to their conformations in the apo form, which is achieved through superimposition of the Fnш-1 domain in the two states using VMD [22]. The two structures shown in **Fig. S3C** are taken as a reference because half of the conformation in their structures is similar to the structure in the intermediate conformation (shown in **Fig. S3B**), *i.e.*, the Fnш-1 domain makes contacts with the L2’ domain of the partner. In the original experimental structure [13] (PDB code: 5KQV), two antibodies occupy the two insulin T2BSs. After removal of the antibodies, two T2BSs on two Fnш-1 domains are exposed and appear to be capable of insulin binding. Because-two insulin T2BSs are exposed in the auto-inhibited conformation while insulin T1BSs are partially or completely buried, it is possible that two insulin molecules may bind to two T2BSs during the auto-inhibited state, which drives the conformation transition from the initial auto-inhibited state to the possible intermediate state(s). Therefore, two insulins are added to two T2BSs on the two Fnш-1 domains according to the T2BS structure as shown in **Fig. S3F** [17, 18]. Superimposition of the two Fnш-1 domains with/without an insulin at each T2BS using VMD [22] helps place the corresponding insulin molecules inside the T2BSs in the intermediate conformation.

The modeled intermediate conformation is symmetric (**Fig. 2B**), and the main feature of this symmetric intermediate state is that there are two insulins bound inside the two T2BSs plus two insulins inside two incomplete T1BSs on the L1 domains, while the Fnш-1 domain makes contacts with the L2’ domain of the partner. The insulin at the incomplete T1BS on the L1 domain is just over against the Fnш-2’ domain of the partner. The αCT’, ID-α’ and ID-β’ domains of the partner are located beside insulin, at the lower and upper inclined sides of insulin, respectively, which appears to make preparation for forming the complete T1BS. This structural model may shed some light on the dynamic process leading to the formation of the complete T1BS. From a different viewing angel (**Fig. S4A**), the intermediate state also appears to be a U-shaped conformation, which might be the reason that it was considered as a possible initial state in some of the earlier IR activation models [17, 18].

#### Construction of the asymmetric active state of IR

In 2018, Lawrence *et al.* determined the cryo-EM structures of the ectodomain in a signaling conformation [15] (PDB code: 6HN4, 6HN5; **Fig. S3C**). Prior to this work, Scapin *et al.* also reported a cryo-EM structure of the ectodomain in a similar conformation [16] (PDB code: 6CE7; **Fig. S3D**). Because only one complete T1BS is bound with insulin [15, 16] (**Fig. S3C** and **S3D**), the full-length IR presents as an asymmetric conformation which is similar to the II-shaped conformation as reported earlier [21]. The main difference between the two asymmetric II-shaped conformations is that the L1’ domains without insulins at the incomplete T1BSs locate over against the Fnш-2 (left panel of **Fig. S3C**) and Fnш-1 (**Fig. S3D**) domains of the partner, respectively [15, 16]. In the present modeling study, these two structures are named as asymmetric state 1 (**Fig. S3C**) and asymmetric state 2 (**Fig. S3D**), respectively, for convenience of discussion.

In order to acquire the complete ectodomain in asymmetric state 2, the missing parts of the ectodomain (**Fig. S3D**) are appended according to the structure in asymmetric state 1 (left panel of **Fig. S3C**), through superimposition of the Fnш-2’ domain. Because the αCT domain near the incomplete T1BS without insulin is missing in the experimental structure, it is necessary to add the αCT domain back into the structure. To do so, the αCT domain is placed into the incomplete T1BS according to its relative position to the L1’ domain of the partner as shown in the complete T1BS, which is accomplished through superimposition of the two L1 domains involved in the complete and incomplete binding sites using VMD [22]. The other missing segments near the T1BSs such as ID-α and ID-β are constructed using SWISS-MODEL [23]. The distance between the two Fnш-3 domains is sufficiently close for dimerization of the tyrosine kinase domains (left panel of **Fig. S3C**). It is predicted that the asymmetric II-shaped conformations likely are in the active states. In line with this prediction, an earlier study by Gutmann *et al.* also reported that the full-length IR in single lipid nanodisc can present an II-shaped conformation in the presence of insulins (**Fig. S1**) and one insulin can induce the conformation transitions [21].

Because the structure of the functional dimer of tyrosine kinase domains is not available in the Protein Data Bank [27], the symmetric functional dimer is constructed according to the available single tyrosine kinase domain [20] (PDB code: 4XLV, right panel of **Fig. S3G**). The juxtamembrane domain makes interactions with a helix in tyrosine kinase domain of the partner [20], which is considered the main structural restraint during the construction of the symmetric dimer. Based on the constructed structure of tyrosine kinase and juxtamembrane dimer, the predicted structure of βCT domain and the available experimental structure of transmembrane domain [19], a new homodimer is modeled by assembling the transmembrane and juxtamembrane domains, the dimer of tyrosine kinase domains, and the βCT domains through superimposition of the common parts with VMD [22]. Then, the new homodimer is appended to the ectodomain in different states. Additionally, the missing insulin(s) at the exposed T2BSs on Fnш-1 domains is (are) added according to the structure of insulin T2BSs. The full-length asymmetric II-shaped conformations corresponding to the structures in **Fig. S3C** and **S3D** are shown in **Fig. 2C** and **Fig. S4B**, respectively. The main difference of the two asymmetric active states is that only one Fnш-1 domain makes contacts with the L2’ domain of the partner. Fnш-1’ domain of the partner makes contacts with the insulin at the complete T1BS, with another insulin at the T2BS. Another T2BS on the Fnш-1 domain is also bound with an insulin in the asymmetric active state 1 (**Fig. 2C**), but in the case of asymmetric active state 2, this same binding site appears to be hindered by the L1’ domain of the partner (**Fig. S4B**). The two tyrosine domains form a functional dimer at their active states. Because the overall conformations of these two asymmetric active states are highly similar, only one conformation (**Fig. 2C**) is shown to represent these two similar states.

#### Construction of the symmetric active state of IR

In the work of Scapin *et al.* [16], there are two other cryo-EM structures of the ectodomain (PDB code: 6CE9 and 6CEB; **Fig. S3E**). Recently, Gutmann *et al.* and Bai *et al.* reported the cryo-EM structures of the complete ectodomain with four insulin molecules bound inside the binding sites [17, 18] (PDB code: 6PXV, 6SOF; **Fig. S3F**). As shown in **Fig. S3E** and **S2F**, the binding of insulins inside the two complete T1BSs appears to result in a symmetric T-shaped conformation, which is also commonly known as an active state [18]. The experimental structures of the ectodomain (**Fig. S3F**) in this conformation are more complete.

To assemble the full-length IR in its symmetric active state, the missing segments or residues in the ectodomain are constructed using SWISS-MODEL [23], and the other domains are appended to the ectodomain using VMD [22] according to the corresponding structures in the full-length II-shaped conformations (**Fig. 2C**). The full-length symmetric active conformation (**Fig. 2D**) is the final state of IR activation. The main characteristics of the symmetric active state are that the two complete T1BSs are on the shoulders of the structures, and two insulin molecules at the two T2BSs appear to be only loosely bound and ready to dissociate from the Fnш-1 domains (**Fig. S3E**).

In summary, according to the above analysis and also based on available experimental structures, it is speculated that the full-length IR might have four representative conformations, which correspond to the four different states of the receptor, namely, the apo form (auto-inhibited state; **Fig. 2A**), the symmetric intermediate state (**Fig. 2B**), the asymmetric active state (**Fig. 2C**), and the symmetric active state (**Fig. 2D**). In total, there are one representative auto-inhibited state, one representative intermediate state, plus two representative active states. It is speculated that the association and disassociation of the insulin molecules into the dynamic binding sites of IR drive the sequential processes leading to the formation of these states.

Here it is of note that the two transmembrane helices in all conformations are confined in two membrane planes by rotation of the transmembrane domains and other related domains using VMD [22]. As there are missing residues in insulin structure, it is replaced with a complete structure (1.60 Å resolution; PDB code: 3I3Z; **Fig. S3H**) [28] through superimposition. Energy minimization of the IR conformations is carried out to ensure that the local structures are reasonable. Backbones of all the conformations are extracted, new side chains are added to the structures with CHARMM-GUI [29], and energy minimization is conducted with NAMD [30].

To visualize the structural changes of the L1∼CR∼L2∼Fnш-1 domains, top views of the four representative conformations of the full-length IR are shown in **Fig. 3**. Based on the modeled structures, it appears that in the apo form, no insulin is present (**Fig. 3A**); in the symmetric intermediate state, two insulins are present at the two incomplete T1BSs plus two insulins at the two T2BSs (**Fig. 3B**); in the asymmetric active state, one insulin is present at the complete T1BS plus two insulins at the T2BSs (**Fig. 3C**); in the symmetric active state, two insulins are present at the two complete T1BSs plus two insulins at the two T2BS (**Fig. 3D**). In the symmetric active state, insulins at the T2BSs can dissociate from the IR, which leads to only two insulins in the conformation as shown in **Fig. S3E**. There is a structural rearrangement of the four domains from the apo form to the symmetric intermediate state. The global and local conformational transitions are discussed separately in more detail below.

**Figure 3.**
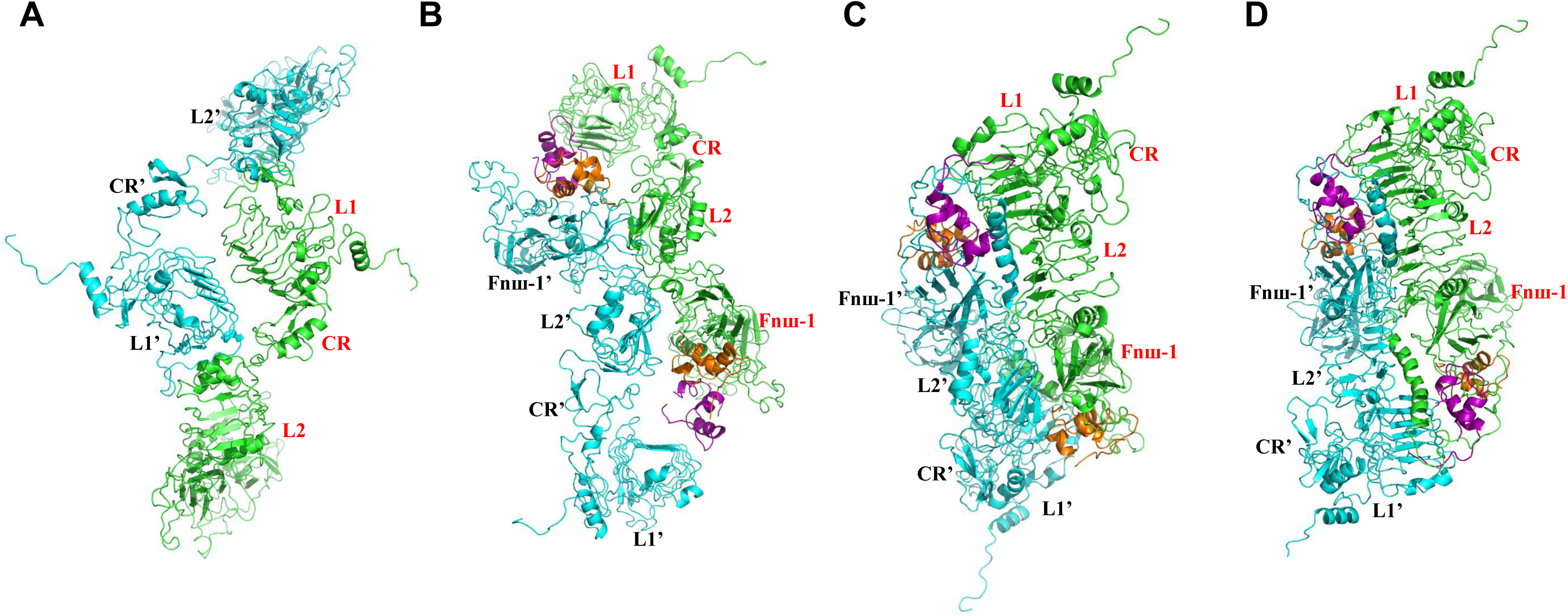
Top view of the four conformations of the full-length IR. **A.** The apo form (auto-inhibited state). **B.** Symmetric intermediate state. **C.** Asymmetric active state. **D.** Symmetric active state. The first 850 residues of IR plus insulin are shown to illustrate the structural changes of the ectodomain. Insulins at the incomplete or complete T1BSs are colored in purple, and insulins at the T2BSs colored in orange.

### Conformational transition pathways

According to the above-described four representative conformations of the full-length IR, the conformational transition pathways can be divided into three steps. The first step (step 1) is from the auto-inhibited state (**Fig. 4A**) to the symmetric intermediate state (**Fig. 4B**), the second step (step 2) is from the symmetric intermediate state (**Fig. 4B**) to the asymmetric or symmetric active states (**Fig. 4C, 4D**), and the third step (step 3) is from the active states (**Fig. 4C, 4D**) back to the auto-inhibited state (**Fig. 4A**). Note that only the overall conformational transitions at this step will be discussed here, and details of the structural changes, especially the structural changes at the insulin binding sites, will be discussed later.

**Figure 4.**
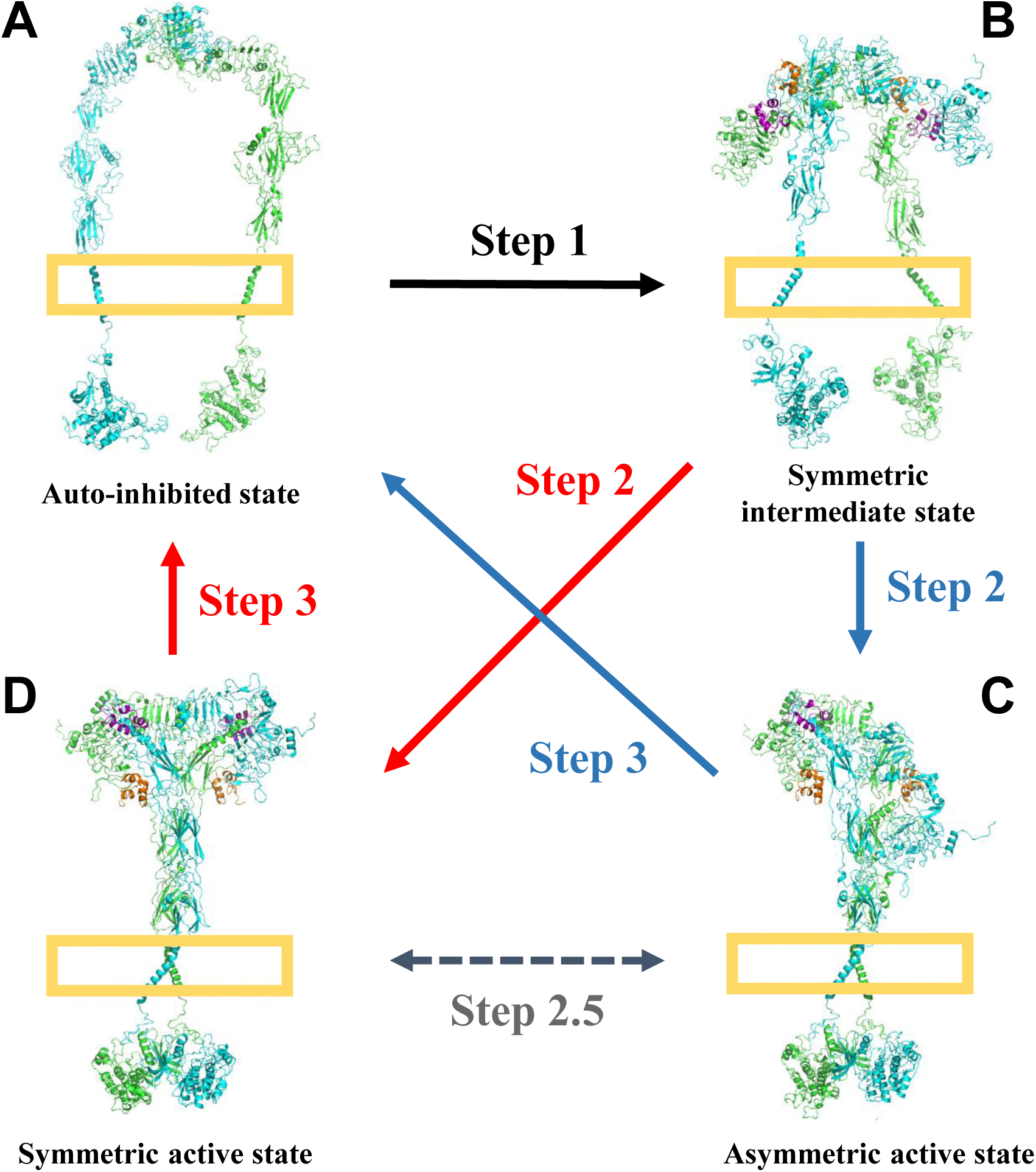
Conformational transition pathways during the process of the full-length IR activation. **A.** The apo form (auto-inhibited state). **B.** Symmetric intermediate state. **C.** Asymmetric active state. **D.** Symmetric active state. The two chains of IRs are colored in green and cyan. Insulins at the incomplete or complete T1BSs are colored in purple, and insulins at the T2BSs colored in orange. The complete conformational change process is artificially divided into the following steps for convenience of description and understanding: **Step 1:** Binding of one or two insulin(s) to its T2BSs would induce major conformational changes from the auto-inhibited state to symmetric intermediate state. In the process, the binding of one or two insulin(s) to the exposed incomplete T1BSs may also facilitate the conformational transitions. **Step 2**: Insulin(s) at the incomplete T1BSs would induce the transition from symmetric intermediate state to asymmetric or symmetric active states. **Step 3**: Dissociation of insulins would induce conformational transition from the asymmetric or symmetric active state back to the auto-inhibited state. Additionally, it is suggested that the two representative conformations at the asymmetric and symmetric active states can be converted via formation or disruption of a complete T1BS, and this process is considered as an intermediate step (**Step 2.5**).

At step 1 of the conformational transition pathway, insulin binding to T2BS may induce the separation of the L1∼CR∼L2∼Fnш-1∼L1’∼CR’∼L2’∼Fnш-1’ dimer (**Fig. 5B, 5C**, **Fig. 6B, 6C**). Then, two insulins can bind to two incomplete T1BSs (**Fig. 5D**, **Fig. 6D**, **Fig. S3B**). Finally, relative movements between L1∼CR∼L2∼Fnш-1 and L1’∼CR’∼L2’∼Fnш-1’ of the partner result in the formation of symmetric intermediate conformation in which the Fnш-1 domain makes contacts with the L2’ domain of the partner (**Fig. 5E**, **Fig. 6E**). The order of insulin binding to the incomplete T1BS and the relative movements between L1∼CR∼L2∼Fnш-1 and L1’∼CR’∼L2’∼Fnш-1’ of the partner are likely to be interchangeable.

**Figure 5.**
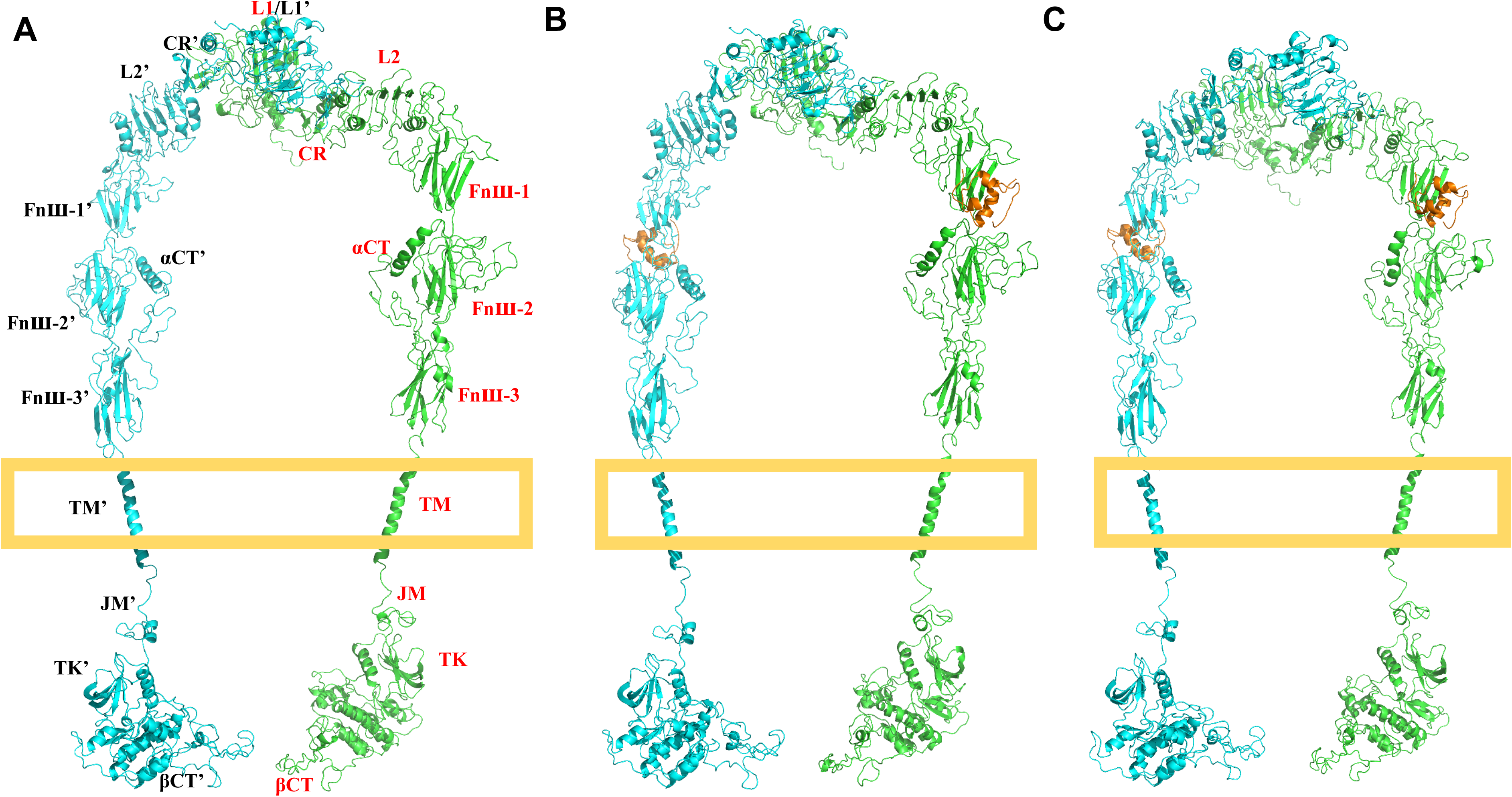
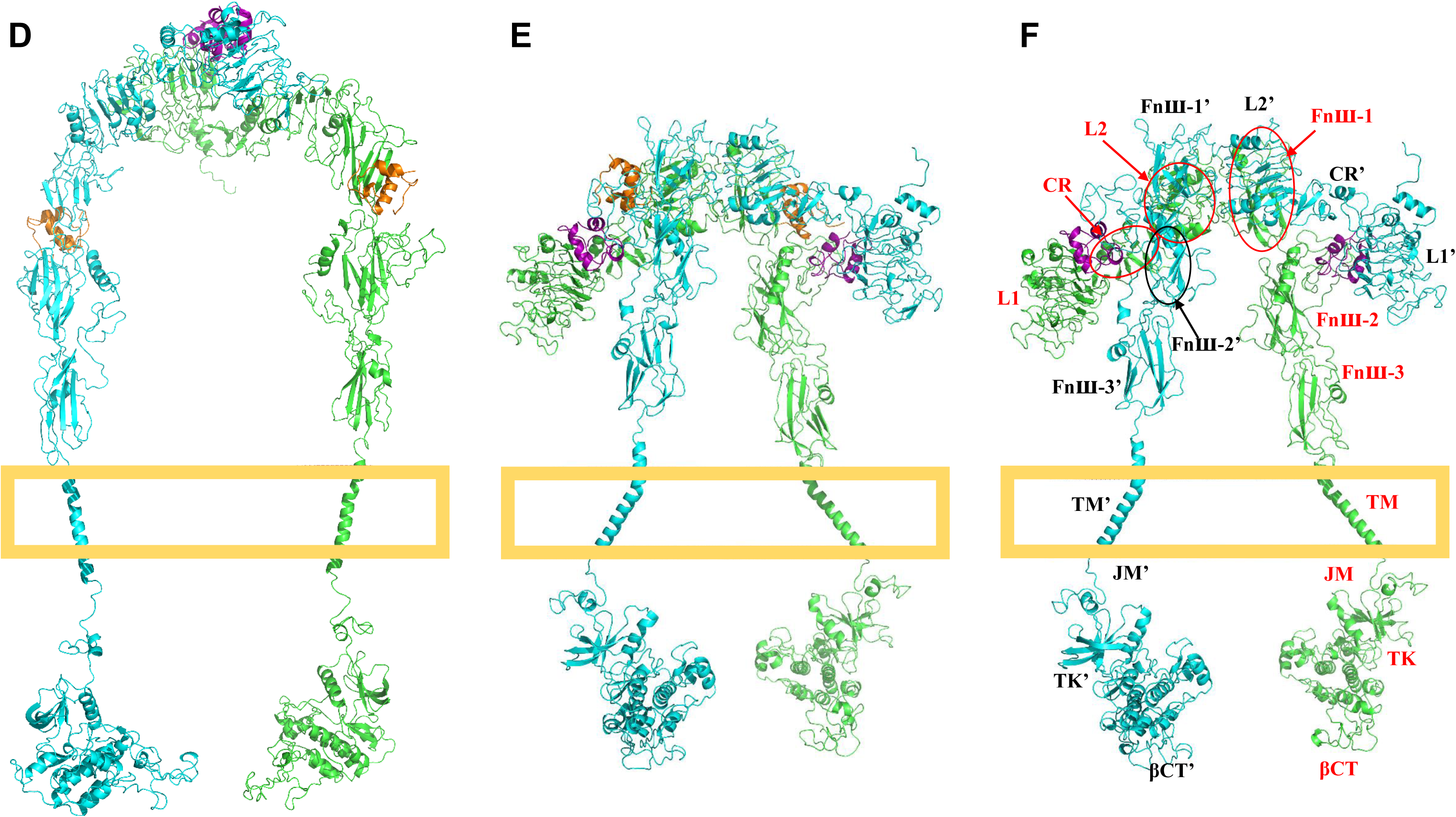
Conformational transition pathway from the apo form (auto-inhibited state) to symmetric intermediate sate. The overall pathway is from **A** to **E** or **F**, and the order of **D** and **E** may be exchanged (or reversed?). **A.** The apo form (auto-inhibited state). **B.** Conformation generated by binding of two insulins to two T2BSs. **C.** Conformation generated through structural rotation. **D.** Conformation generated by binding of two insulins to the incomplete T1BSs. **E.** Symmetric intermediate state (with four insulins). **F.** Symmetric intermediate state (with two insulins at the two incomplete T1BSs). The two chains of IRs are colored in green and cyan. The insulins at the incomplete or complete T1BSs are colored in purple, and insulins at the T2BSs colored in orange.

**Figure 6.**
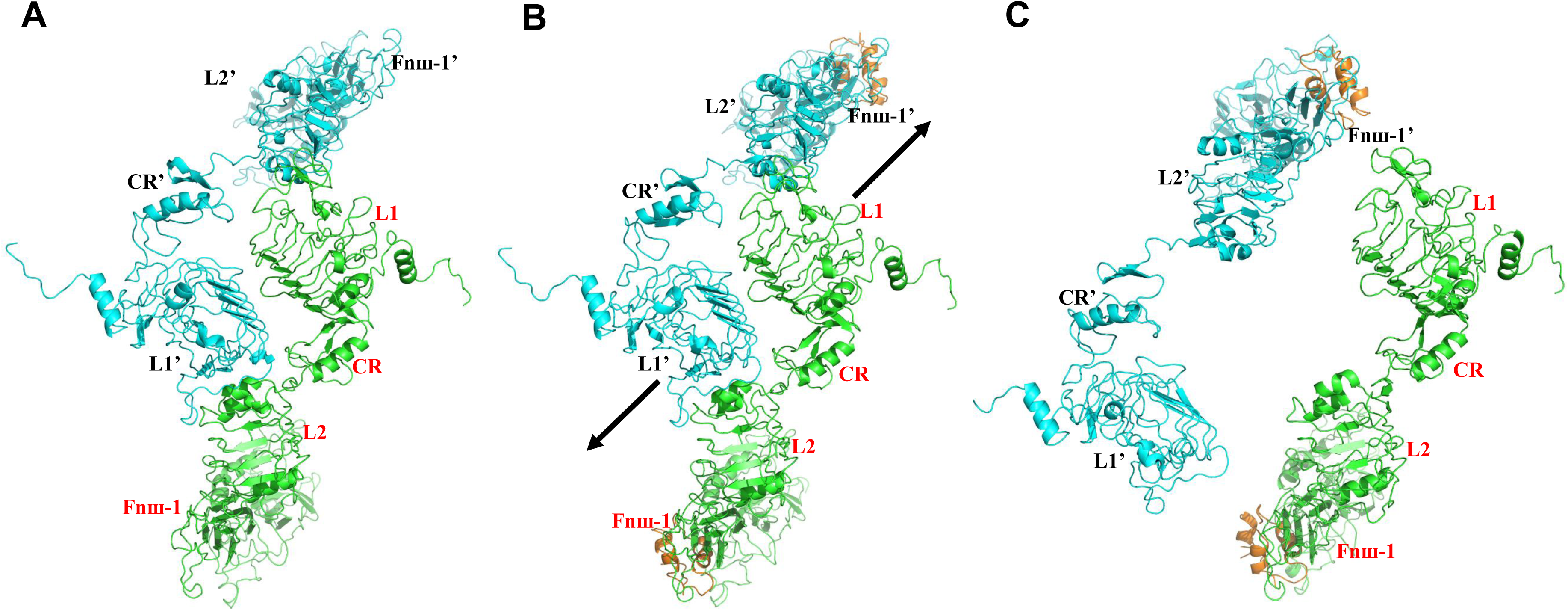
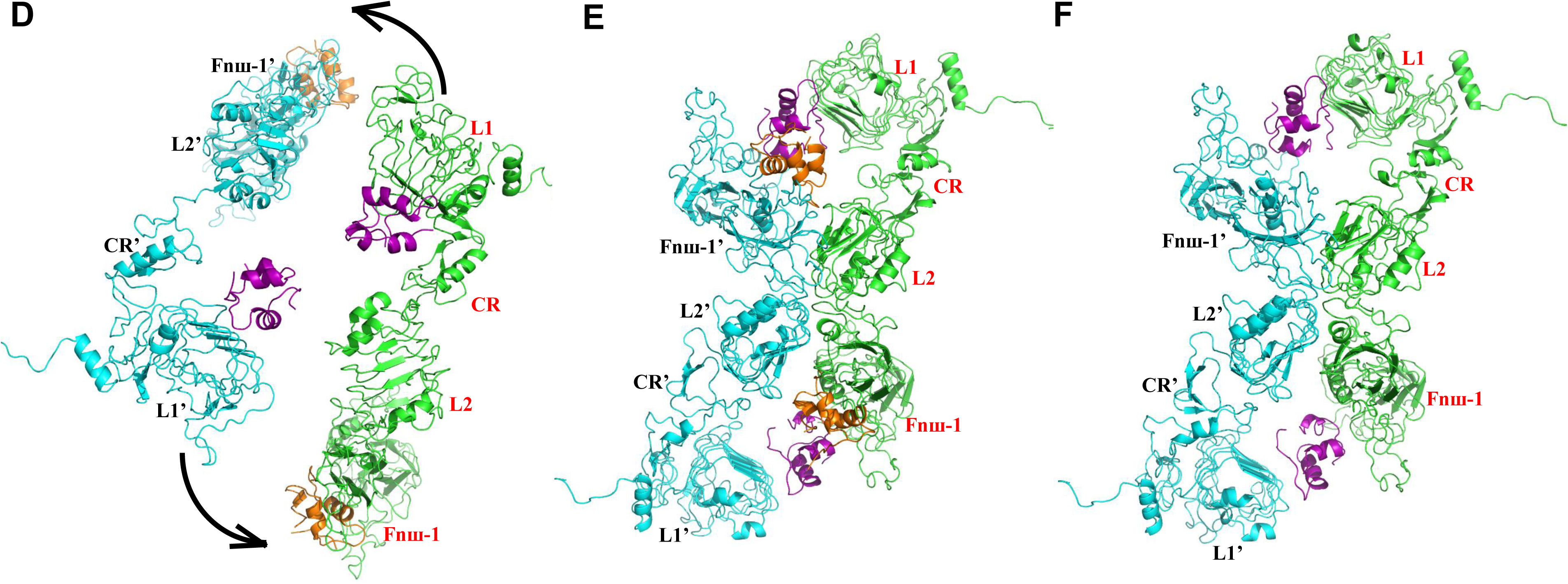
Top view of structural arrangements of the L1∼CR∼L2∼Fnш-1∼L1’∼CR’∼L2’∼Fnш-1’ dimer during conformational change from the apo form (auto-inhibited state) to symmetric intermediate sate. The pathway is from **A** to **E** or **F**, and the order of **D** and **E** may be exchanged. **A.** The apo form (auto-inhibited state). **B.** Conformation generated by binding of two insulins to the two T2BSs. **C.** Conformation generated through structural rotation. **D.** Conformation generated by binding of two insulins to the two incomplete T1BSs. **E.** Symmetric intermediate state (with four insulins). **F.** Symmetric intermediate state (with two insulins at the two incomplete T1BSs). The two chains of IRs are colored in green and cyan. The insulins at the incomplete or complete T1BSs are colored in purple, and insulins at the T2BSs colored in orange.

Step 2 of the pathway has two different manifestations. If only one complete T1BS is formed (**Fig. 4C**), the conformational change would be asymmetric. Gutmann *et al.* have proposed that binding of one ligand is sufficient to induce the transition of the IR ectodomain from the U-shaped conformation to the T-shaped conformation, based on available experimental data [21]. Therefore, it is suggested that the binding of an insulin molecule at one complete T1BS likely would be sufficient to initiate the process of IR activation (**Fig. 4C**). If the formation of two complete T1BSs is accomplished simultaneously, then the conformational transitions would be symmetric (**Fig. 4B, 4D**), and the functional tyrosine kinase dimer likely would also be formed during this process.

Additionally, as an intermediate step (step 2.5), there may exist reversible conformational transitions between the asymmetric (**Fig. 4C**) and symmetric (**Fig. 4D**) active states. Formation of the other complete T1BS in the asymmetric active conformation (**Fig. 4C**) would make the IR structure symmetric again (**Fig. 4D**). As a final step (step 3), it is speculated that dissociation of the insulin molecules from the IR would induce the conformational transition from the asymmetric or symmetric active states (**Fig. 4C, 4D**) back to the initial auto-inhibited state (**Fig. 4A**).

Lastly, it is of note that because the intermediate conformation of the full-length IR bound with insulins was observed when the two legs of the transmembrane part are individually inserted into two lipid nanodiscs [21], it is assumed that the same (or very similar) intermediate conformation, likely in a transient state, would also be formed with one nanodisc. This intermediate state may exist in the process at step 3 with a low probability. It is speculated that the formation of the complete T1BS from its incomplete precursor may be a facile process, and the intermediate state (as 2T-shaped conformation in **Fig. 1**) may only exist in the process of conformational activation from the initial auto-inhibited state to the active states (but may not exist in the opposite process).

### Details of the structural changes and motions of different domains in the full-length IR

#### Analysis of Step 1

To probe the structural changes in different domains during the process of conformational transitions, the structures between the same domains in each pair of conformations shown in **Fig. 2** and **Fig. S4B** are superimposed using VMD [22], and the mean RMSDs between the Cα atoms in the same domains are calculated and compared (results are summarized in **Table 1**). The mean RMSD values are >4.0 Å for ID-α, αCT, ID-β, TM and JM, which reflect the high degree of flexibility and large structural changes and motions in these domains. The ID-α∼αCT∼ID-β domains form a long loop, and structural changes in this long loop are important to form the complete T1BSs. TM is a helix, and the flexibility of TM and JM domains is mainly in the JM domain. Rigid motions are dominant for other domains during the process.

**Table 1.**
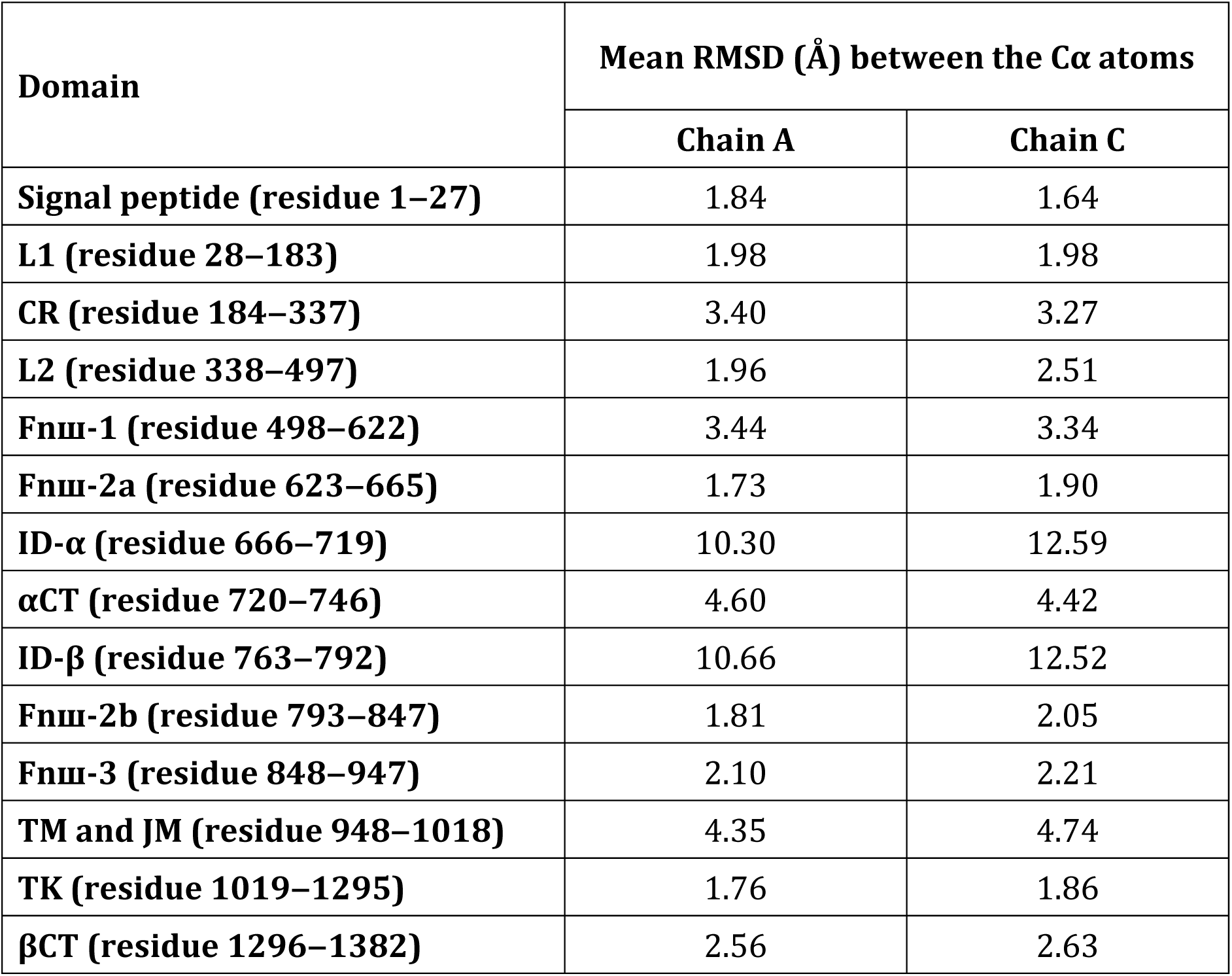
Mean RMSD (Å) between the Cα atoms of different domains in the representative IR conformations shown in Fig. 2 and Fig. S4B.

Structural rearrangement of IR’s L1∼CR∼L2∼Fnш-1 domains from IR’s apo form to its symmetric intermediate state (**Fig. 6**) has been briefly discussed above. The L1∼CR domains make contacts with the L2’ domain of the partner in the apo form (**Fig. 6A**). After structural rearrangement, the L2 domain makes interactions with the Fnш-1’ domain of the partner (**Fig. 6E, 6F**) in the symmetric intermediate state.

More details of the structural rearrangement are investigated using Gaussian network model (GNM) [31, 32]. The fluctuations of residues in the L1∼CR∼L2∼L1’∼CR’∼L2’ dimer (PDB code: 2HR7) and the etodomain at active state (PDB code: 6PXV) are calculated first based on GNM, and the correlation coefficient between the theoretical B-factors derived from GNM and the corresponding experimental B-factors in the L1∼CR∼L2∼L1’∼CR’∼L2’ dimer (PDB code: 2HR7) and the ectodomain (PDB code: 6PXV) is approximately 0.67 and 0.83, respectively (**Fig. S5A** and **S5B**), which illustrates the applicability of GNM in IR.

The ectodomain of IR’s auto-inhibited conformation with two insulins bound to the T2BSs (**Fig. 7A**) is extracted for simulating the disassociation of the L1∼CR∼L2∼L1’∼CR’∼L2’ dimer. There are 52 inter-residue contacts at the interface of the L1∼CR∼L2∼L1’∼CR’∼L2’ dimer (cutoff = 7 Å, and each residue is represented by the Cα atom), and the interface contacts can be categorized into four regions (**Fig. 7B**). The first region includes only one inter-residue contact between L1 and CR’ domains, the second one contains four inter-residue contacts between L1 and L1’ domains, the third one involves ten contacts between L2’ domain and the partner chain and one contact between CR and CR’ domains, and the fourth one contains thirty-five contacts between L2 or CR domain and the partner chain (note that contacts in the third region are not counted here).

**Figure 7.**
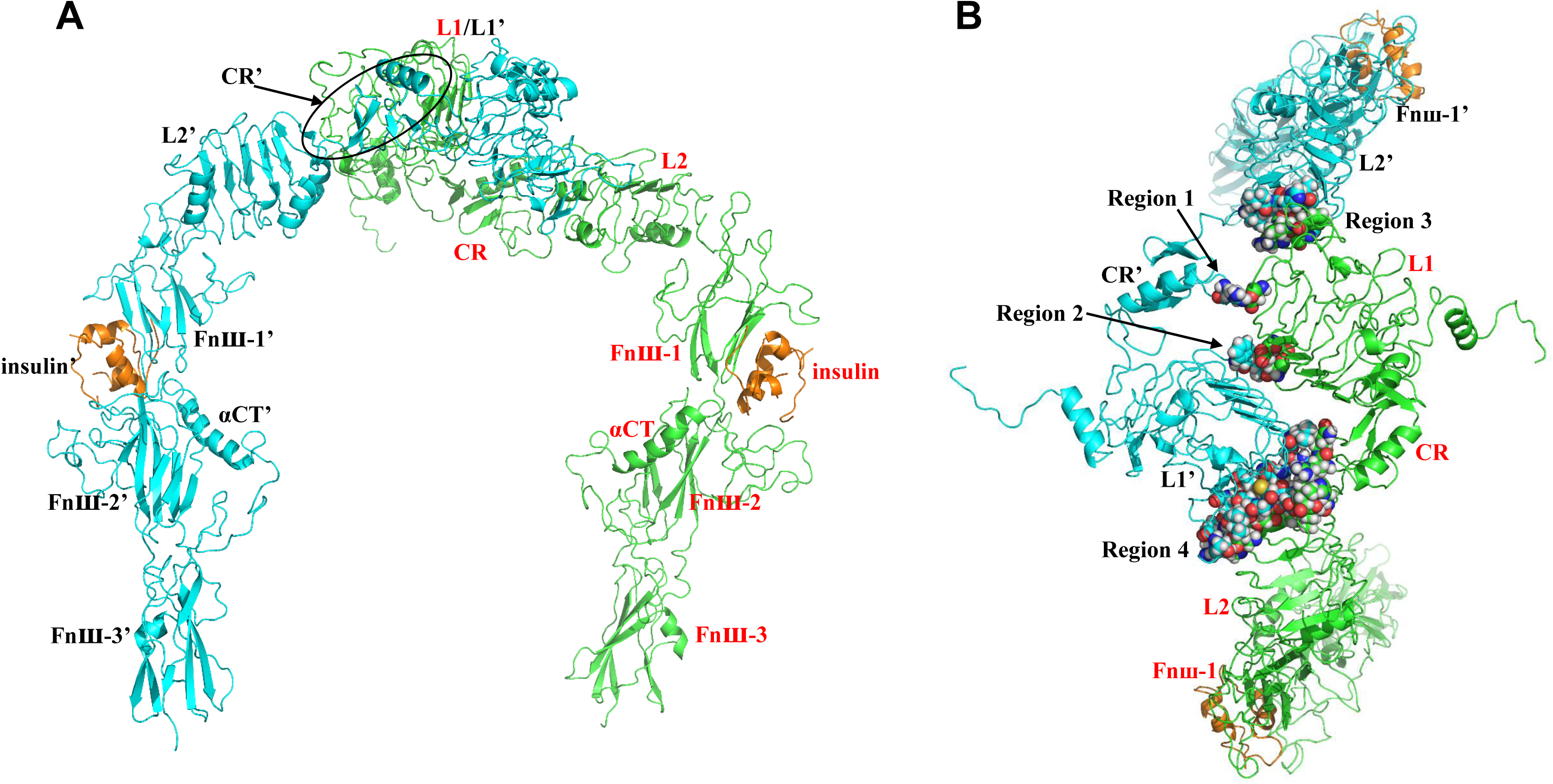
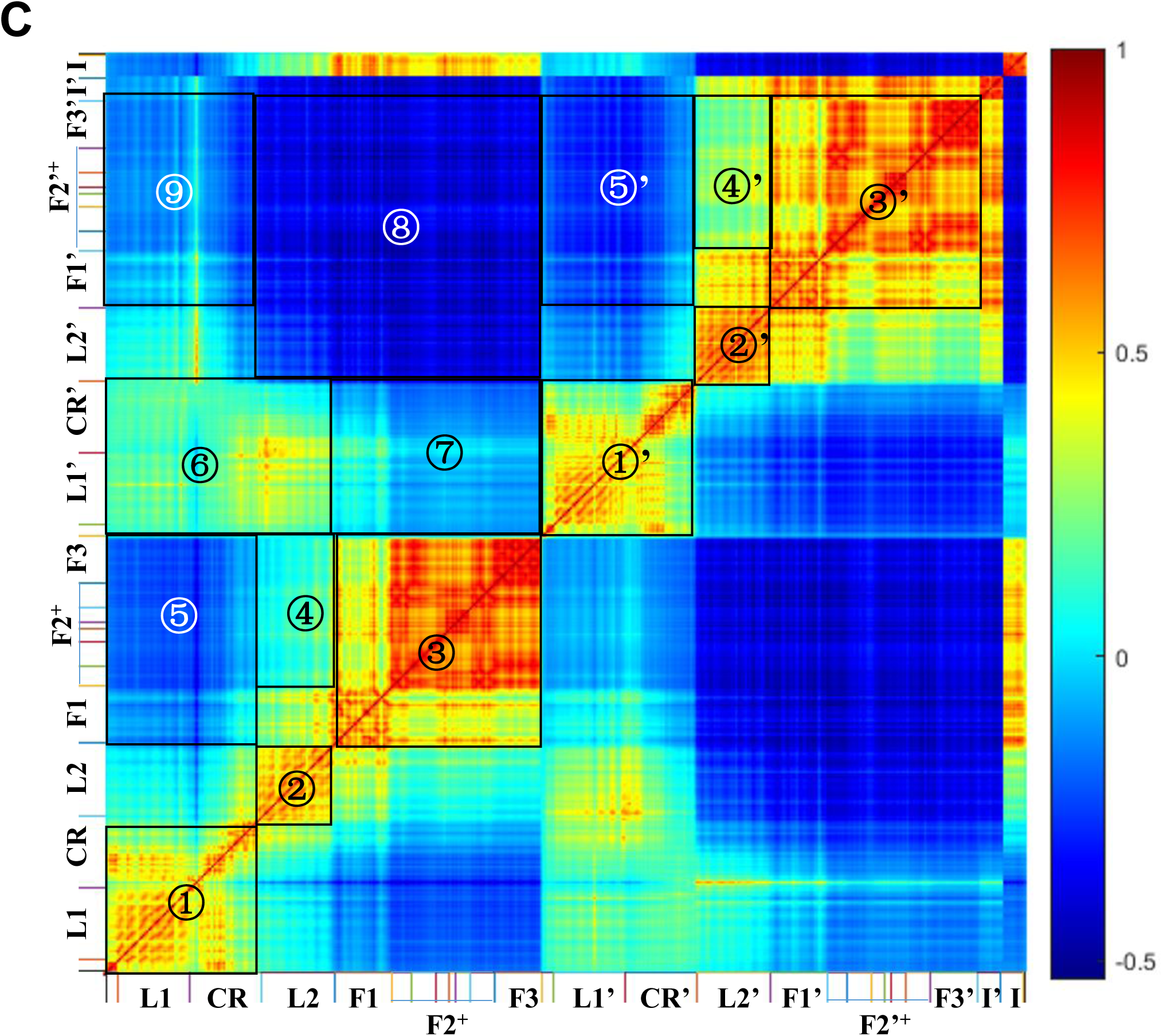
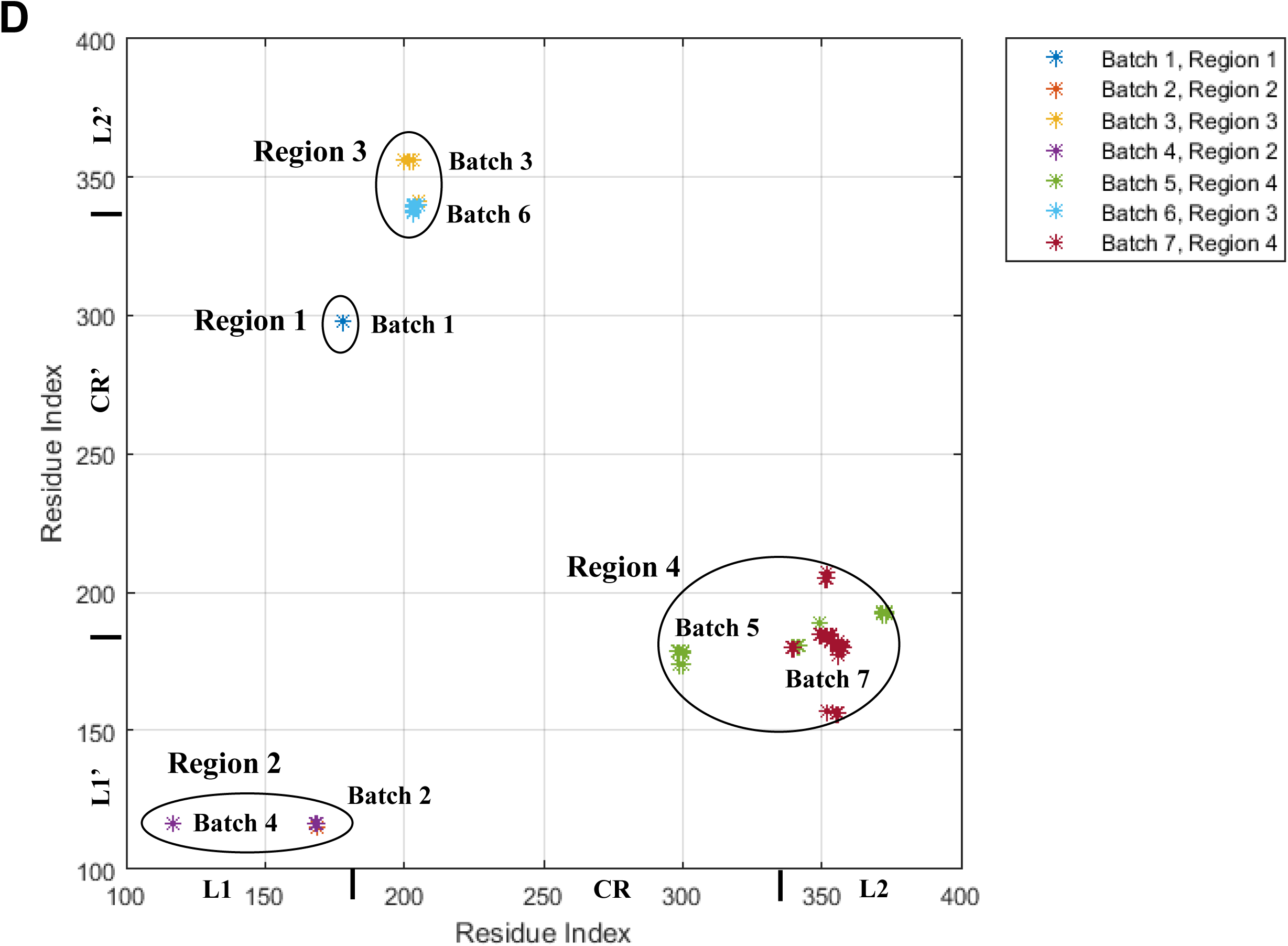
Interface contacts and normalized inter-residue cross-correlation fluctuations of the ectodomain in the auto-inhibited conformation with two insulins at the T2BSs, and the disappearance order of the interface contacts derived from the iterative application of GNM. A. Side view of the ectodomain in the auto-inhibited conformation with two insulins at the two T2BSs on the Fnш-1/1’ domains. **B.** Top view of the interface contacts in the L1∼CR∼L2∼Fnш-1∼L1’∼CR’∼L2’∼Fnш-1’ dimer. The interface contacts can be categorized into four regions: there is one interface contact in the first region, four in the second, eleven in the third, and twenty-six in the fourth. The two chains of IRs are colored in green and cyan. The insulins at the T2BSs colored in orange. **C.** Normalized inter-residue cross-correlation fluctuations of the ectodomain in the auto-inhibited conformation with two insulins derived from GNM. The different domains in IR are labeled along the horizontal and vertical axes. F1, F2 and F3 are the abbreviations of the Fnш-1 domain, the Fnш-2 (Fnш-2a and Fnш-2b) domain and the Fnш-3 domain, respectively. F2^+^ are defined as the structures of Fnш-2a, ID-α, αCT, ID-β and Fnш-2b domains. Different regions in the figure are labeled for convenience of description. **D.** Disappearance order of the interface contacts derived from the iteratively application of GNM. The interface contacts disappear in seven batches, which takes place in these four regions in an alternate manner. The labels along horizontal axis are the residue index and the corresponding domains in one partner chain, and the ones along the vertical axis are the residue index and the corresponding domains in the other partner chain.

The normalized inter-residue cross-correlation fluctuations (NIRCCFs) are calculated to explore the relative movements of the domains in the ectodomain. The values of NIRCCFs range from ‒1 to 1. When the value becomes more positive (closer to 1), the movement directions of the two involved residues would be more similar, and when the value becomes more negative (closer to ‒1), the movement directions of the two involved residues would be more opposite from each other. As shown in **Fig. 7C**, the positive NIRCCFs in the interior of L1∼CR/L1’∼CR’ domains (region ①/①’), L2/L2’ domain (region ②/②’), Fnш-1∼Fnш-2^+^∼Fnш-3/Fnш-1’∼Fnш-2’^+^∼Fnш-3’ (F1∼F2^+^∼F3/F1’∼F2’^+^∼F3’) domains (region ③/③’) indicate that the interior residue-residue interactions in these domains make each of these domains compact. Note that the Fnш-2 (F2) domain is composed of Fnш-2a and Fnш-2b domains, and the Fnш-2^+^ (F2^+^) refers to the complex structure containing Fnш-2a, ID-α, αCT, ID-β and Fnш-2b domains. Because L2/L2’ domains connect with Fnш-1/ Fnш-1’ (F1/F1’) domains directly and Fnш-2^+^∼Fnш-3/Fnш-2’^+^∼Fnш-3’ (F2^+^∼F3/F2’^+^∼F3’) domains indirectly, the L2 domain produces positive NIRCCFs with Fnш-1 domain, but weak NIRCCFs with Fnш-2^+^∼Fnш-3 (F2^+^∼F3) domains (region ④), the L2’ domain yields positive NIRCCFs with the Fnш-1 domain, but weak positive NIRCCFs with Fnш-2^+^∼Fnш-3 (F2^+^∼F3) domains (region ④’). This information implies that the structure is not exactly symmetric. The L1∼CR and L1’∼CR’ domains yield negative NIRCCFs with Fnш-1∼Fnш-2^+^∼Fnш-3 (F1∼F2^+^∼F3) and Fnш-1’∼Fnш-2’+∼Fnш-3’ (F1’∼F2’^+^∼F3’) domains (region ⑤, ⑤’, ⑦, ⑨), L1’∼CR’ domains produce weak positive NIRCCFs with L1∼CR∼L2 domains (region ⑥), and there exist negative NIRCCFs between L2∼Fnш-1∼Fnш-2^+^∼Fnш-3 (L2∼F1∼F2^+^∼F3) domains and L2’∼Fnш-1’∼Fnш-2’+∼Fnш-3’ (L2’∼F1’∼F2’^+^∼F3’) domains (region ⑧), which are consistent with the structural features. Because of the interactions between insulins and T2BSs, insulin and insulin’ (I and I’) yield positive NIRCCFs with Fnш-1∼Fnш-2^+^∼Fnш-3 (F1∼F2^+^∼F3) domains and Fnш-1’∼Fnш-2’^+^∼Fnш-3’ (F1’∼F2’^+^∼F3’) domains, respectively. In general, the intra-residue contacts in the interior of the domains and the inter-residue contacts between different domains would make the movement directions of the corresponding structures very similar. If two different domains are connected by a hinge region, the movement directions of the two domains would be opposite, which can present an open-to-close or close-to-open conformational transitions.

The disappearance order of the interface contacts is predicted according to the distance fluctuations derived from the iterative application of GNM. As shown in **Fig. 7D**, the interface contacts disappear in seven batches, which takes place in four regions alternately. Overall, the interface contacts in the first and second regions disappear first, followed by the disappearance of contacts in the third and fourth regions.

As the disappearance of the interface contacts (**Fig. S6A, S6B** and **S6C**), there are gradual increases in the positive NIRCCFs in the interior of L1∼CR/L1’∼CR’ domains, L2/L2’ domains, and Fnш-1∼Fnш-2^+^∼Fnш-3/Fnш-1’∼Fnш-2’^+^∼Fnш-3’ (F1∼F2^+^∼F3/F1’∼F2’^+^∼F3’) domains. After the loss of the most of the interface contacts, the NIRCCFs between L1∼CR domains and L2’∼Fnш-1’∼Fnш-2’^+^∼Fnш-3’ (L2’∼F1’∼F2’^+^∼F3’) domains become negative, and those between L1’∼CR’ domains and L1∼CR∼L2∼Fnш-1∼Fnш-2^+^∼Fnш-3 (L1∼CR∼L2∼F1∼F2^+^∼F3) domains become weak negative. This information indicates that the movement directions between the two partners are opposite after the disappearance of interface contacts, which facilitate the structural rearrangement of the L1∼CR∼L2∼L1’∼CR’∼L2’ domains.

The process of dissociation of the L1∼CR∼L2 dimer in the auto-inhibited conformation may be also influenced by the other insulins except the ones bound to T2BSs. To further explore the initial structural changes induced by the binding of insulin(s) in the auto-inhibited state, top views of the L1∼CR∼L2∼Fnш-1 dimer bound with insulins in the auto-inhibited state are shown in **Fig. 8**. There are two T2BSs on the Fnш-1/1’ domains in the auto-inhibited state, and both are exposed and ready for insulin binding. As described and discussed above, the insulin binding to the two T2BSs likely is an important driving force for structural rearrangement of the L1∼CR∼L2∼Fnш-1 dimer from the apo form (auto-inhibited state) to the symmetric intermediate sate. Additionally, there appear to be two incomplete T1BSs: while one is partially exposed on the L1 domain (**Fig. 8A**), the other one on the L1’ domain is hidden in the interface of the dimer (**Fig. 8B**). Insulin binding to the partially-exposed incomplete T1BS may induce the initial relative movements between the two L1/L1’ domains, which facilitates the structural rearrangement of the dimer. It should be noted that the asymmetric active state may be also related to the partially-exposed and completely hidden incomplete T1BSs on the L1/L1’ domains in the auto-inhibited state.

**Figure 8.**
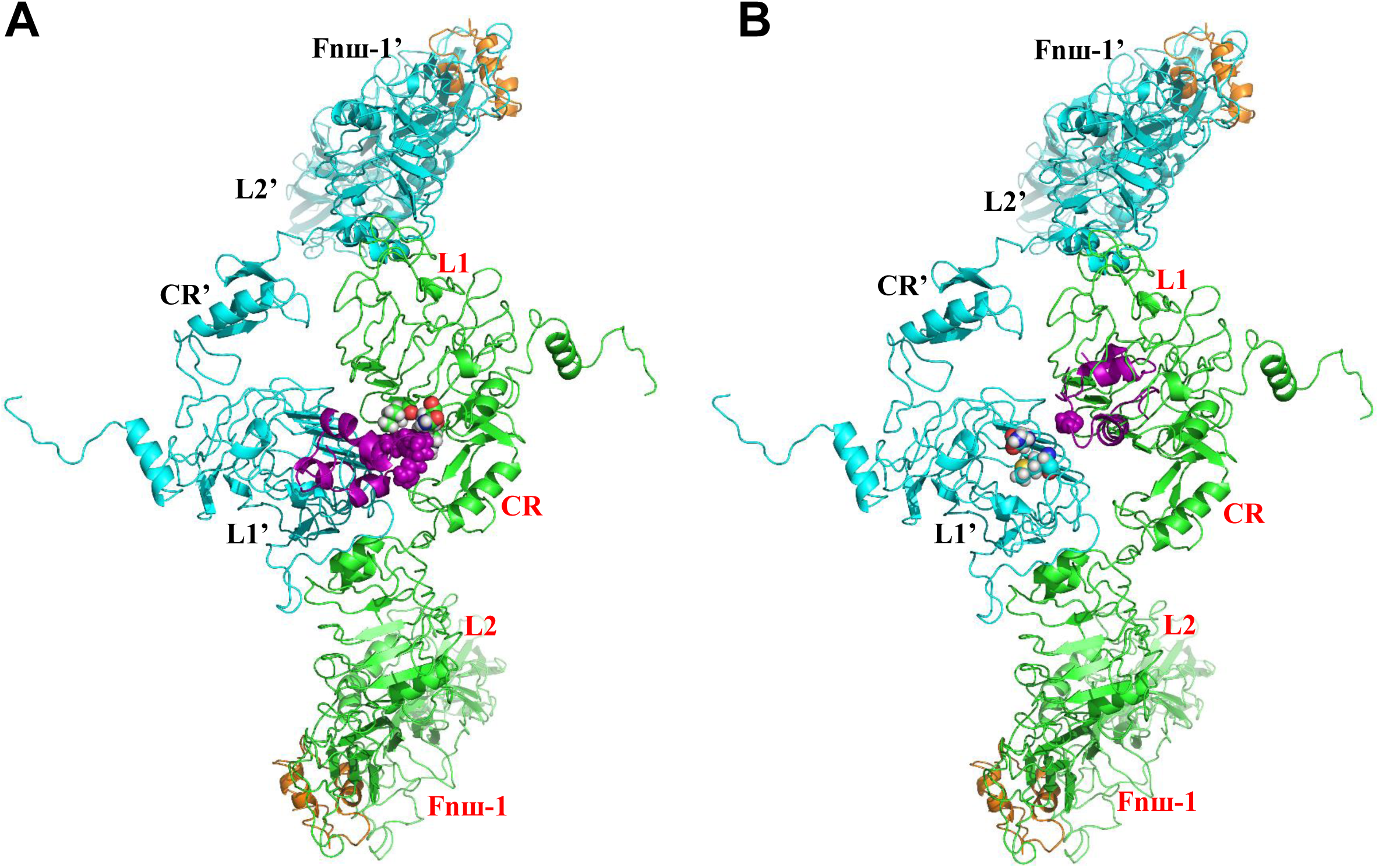
Top view of the L1∼CR∼L2∼Fnш-1∼L1’∼CR’∼L2’∼Fnш-1’ dimer in the auto-inhibited state for exhibition of the primitive insulin binding sites. **A.** Top view of the L1∼CR∼L2∼Fnш-1∼L1’∼CR’∼L2’∼Fnш-1’ dimer with one insulin at the partially-exposed incomplete T1BS on the L1 domain. **B.** Top view of the L1∼CR∼L2∼Fnш-1∼L1’∼CR’∼L2’∼Fnш-1’ dimer with one insulin at the hidden incomplete T1BS on the L1’ domain. The two chains of IRs are colored in green and cyan. The insulins at the incomplete T1BSs are colored in purple, and insulins at the T2BSs colored in orange. The residues involved the interface contacts between the inulin and the incomplete T2BS(s) are shown in sphere.

Moreover, the process for the formation of complete T1BS can also be derived from the intermediate and active states. As shown in **Fig. 9A**, insulin is bound to the L1 domain of an incomplete T1BS, and over against the Fnш-2’ domain of the partner simultaneously. The ID-α’∼αCT’∼ID-β’ domains (a long loop) of the partner are around the insulin. Insulin makes contacts with the L1 and αCT domains of the partner simultaneously. The long loop (*i.e.*, the ID-α’∼αCT’∼ID-β’ domains) and the Fnш-1’ domain of the partner make structural rearrangements. Interaction between the L2 domain and the Fnш-1’ domain of the partner is disrupted, and the L1 and CR domains and the bound insulin move along with the αCT’ domain of the partner, which makes insulin to contact the Fnш-1’ domain of the partner (**Fig. 9B**). In the complete T1BS, insulin makes contacts with the L1 domain as well as the αCT’ and Fnш-1’ domains of the partner.

**Figure 9.**
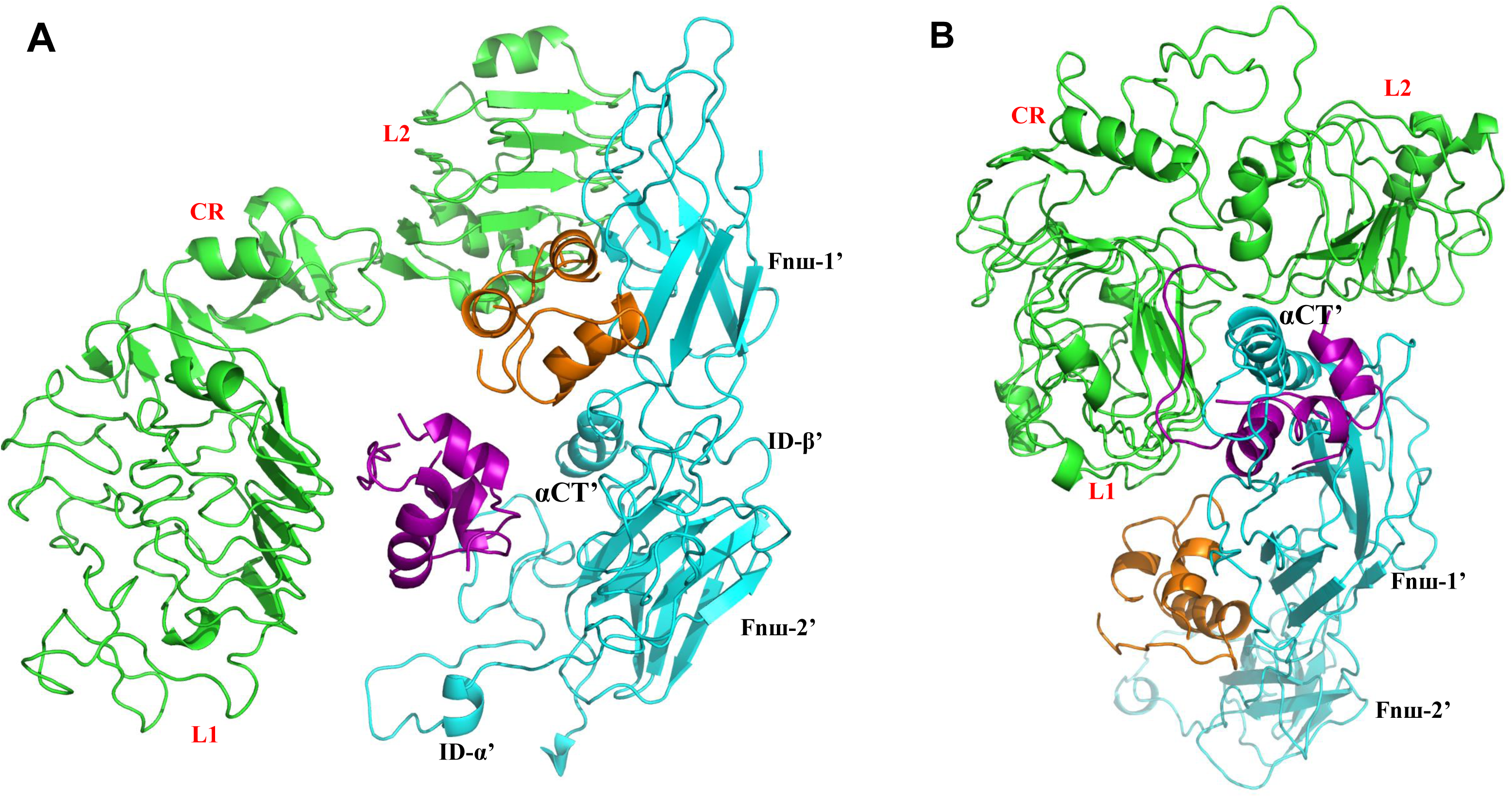
Formation of the complete insulin T1BS. **A.** Incomplete T1BS at intermediate state. **B.** Complete T1BS at the active state. The two chains of IRs are colored in green and cyan. The insulins at the incomplete or complete T1BSs are colored in purple, and insulins at the T2BSs colored in orange.

#### Analysis of Step 2

The ectodomain of the full-length IR in the symmetric intermediate conformation with four insulins (**Fig. 10A**) is extracted for the simulating the disruption of the interactions in the L2∼Fnш-1∼L2’∼Fnш-1’ dimer. In the ectodomain of the symmetric intermediate conformation with four insulins, there are 52 inter-residue contacts at the interface of the L2∼Fnш-1∼L2’∼Fnш-1’ dimer (cutoff = 7 Å, each residue is represented by the Cα atom). According to their positions in the structure (**Fig. 10A**), the interface contacts can be categorized into three regions. The first region (region 1) at the bottom of the dimer incorporate twenty interface contacts. The second and third regions (region 2 and 3) at the top of the dimer contains only one and four inter-residue contacts at the interfaces of L2∼Fnш-1’ domains and L2’∼Fnш-1 domains, respectively. The first regions at the bottom of the dimer can be divided into four sub-regions according to the sequence and structural domains (**Fig. 10A** and **10B**). In the first, second, third and fourth sub-regions (region 1.1, 1.2, 1.3 and 1.4) at the bottom of the dimer, there are ten interface contacts in the L2∼Fnш-1’ domains (region 1.1), two interface contacts in the L2∼L2’ domains (region 1.2), six interface contacts in the L2∼Fnш-1’ domains (region 1.3) and two interface contacts in the Fnш-1∼Fnш-1’ domains (region 1.4), respectively.

**Figure 10.**
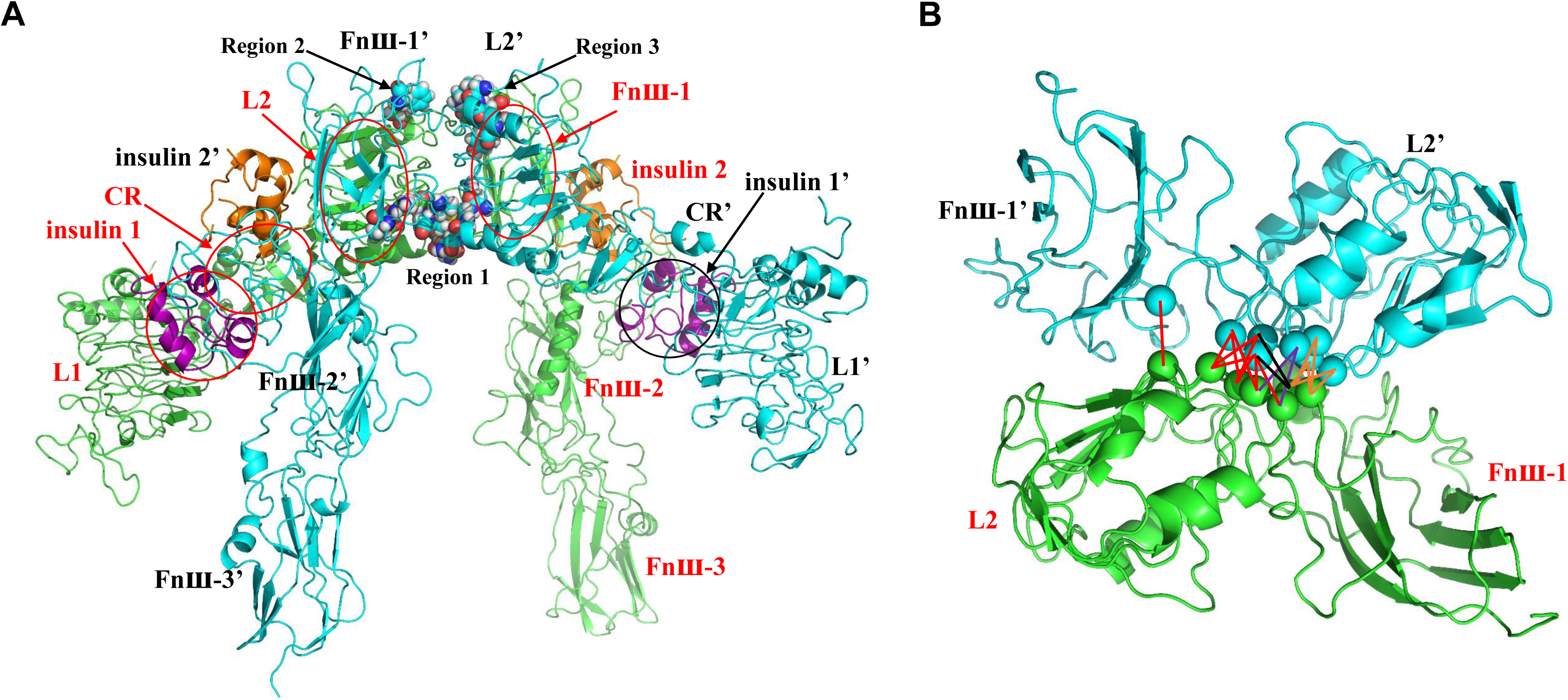
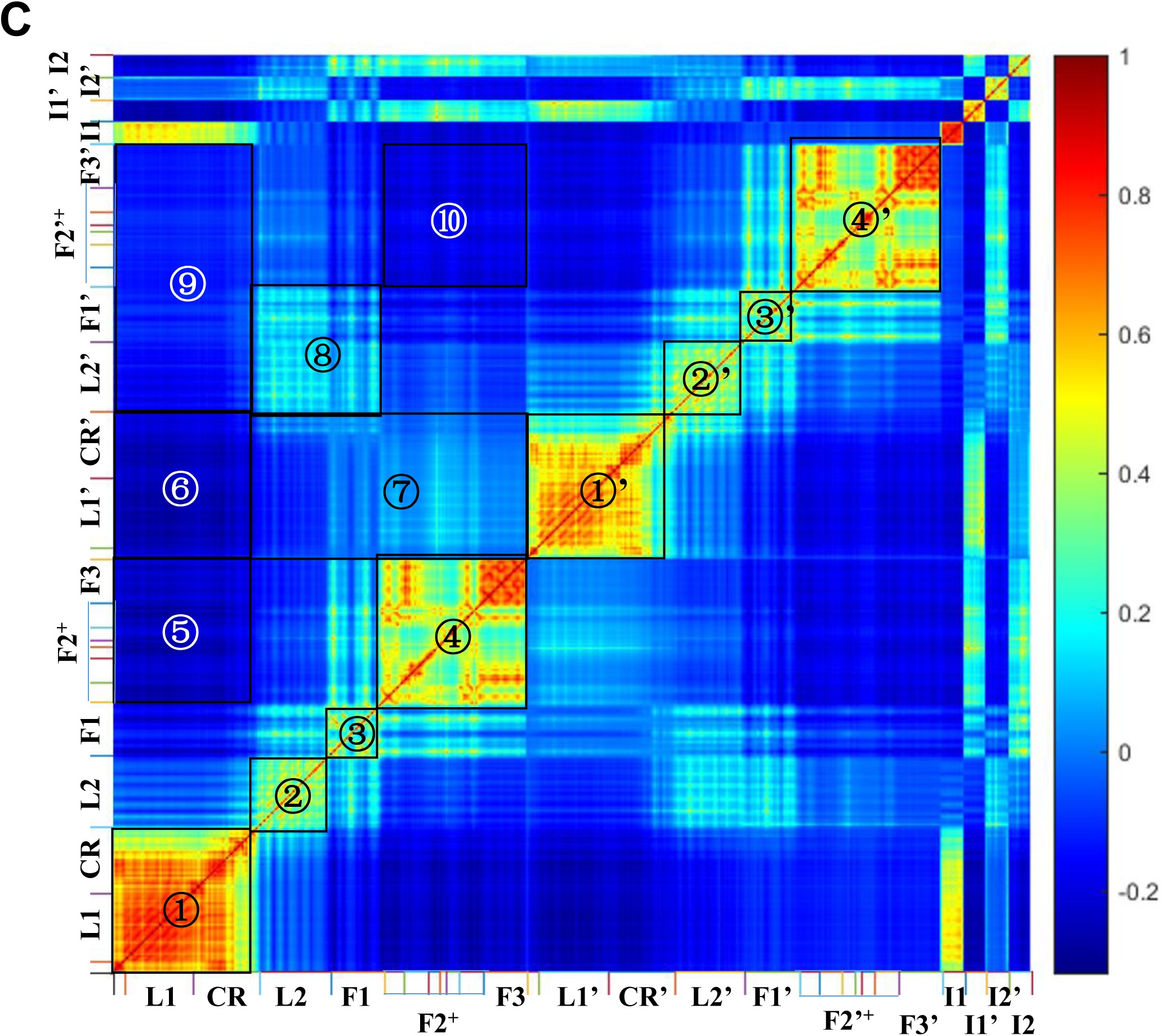
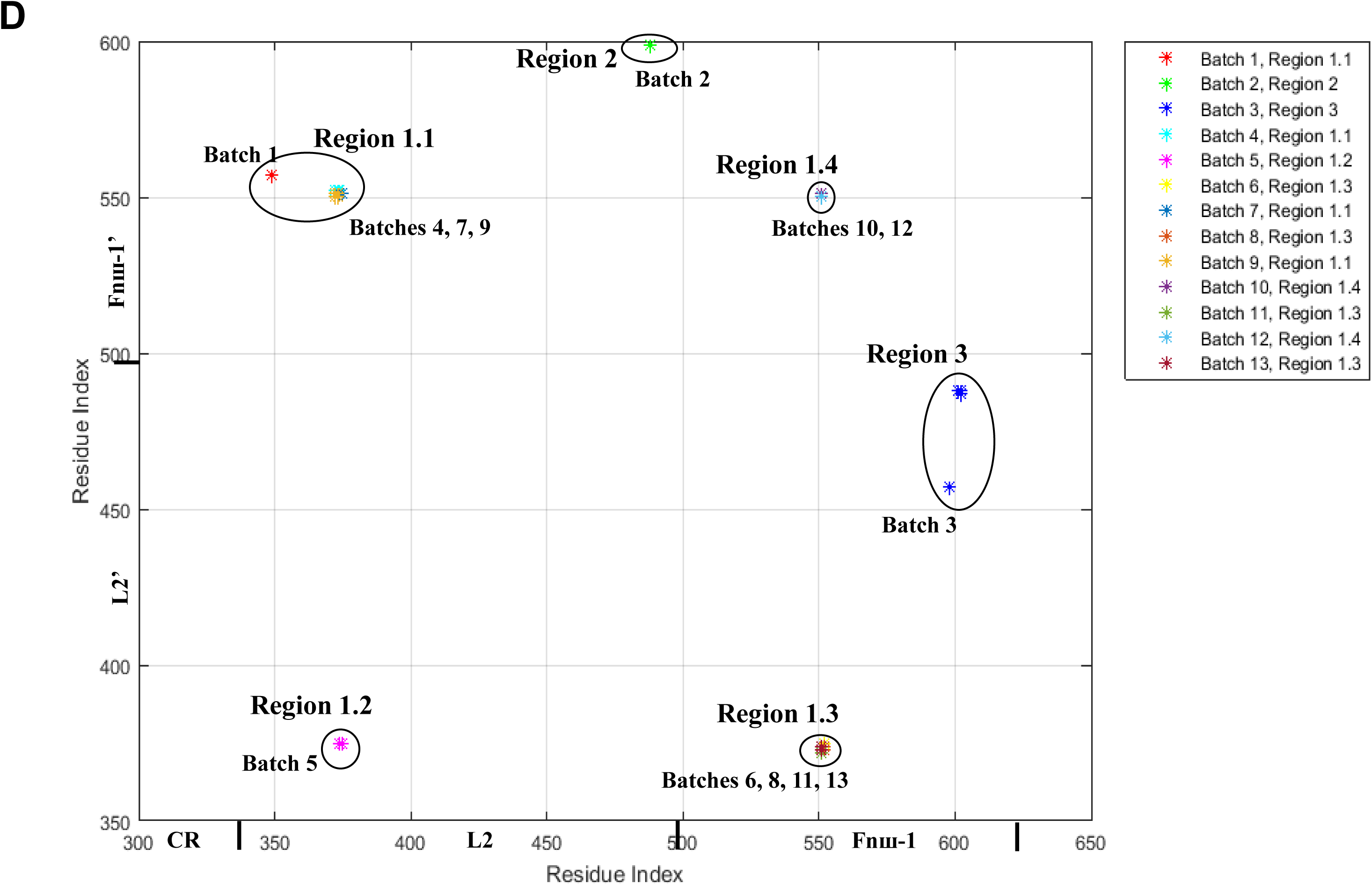
Interface contacts and normalized inter-residue cross-correlation fluctuations of the ectodomain in the symmetric intermediate conformation with four insulins and the disappearance order of the interface contacts derived from the iterative application of GNM. **A.** Side view of the ectodomain in the symmetric intermediate conformation with two insulins at the two T2BSs on the Fnш-1/ Fnш-1’ domains and two insulins at the two incomplete T1BSs on the L1/L1’ domains. There are 25 inter-residue contacts at the interface between the L2∼Fnш-1 and L2’∼Fnш-1’ domains. They can be categorized into three regions. The first region (region 1) locates at the bottom of the dimer, and the second and third regions (region 2 and 3) at the top of the dimer. The two chains of IRs are colored in green and cyan. The insulins at the T2BSs colored in orange. Each pair of arrow and ellipse in red points to a domain in one partner chain which is colored in green; likewise, the pair of arrow and ellipse in black points to a domain in the other partner chain which is colored in cyan. **B.** Bottom view of the inter-residue contacts in region 1 at the interface of the L2∼Fnш-1∼L2’∼Fnш-1’ dimer. The interface contacts can be categorized into four sub-regions/classes. The first, second, third and fourth sub-regions are composed of the inter-residue contacts at the interfaces of L2∼Fnш-1’ domains, L2∼L2’ domains, L2’∼Fnш-1 domains and Fnш-1∼Fnш-1’ domains, respectively, which are colored in red, purple, orange and black, respectively. **C.** Normalized inter-residue cross-correlation fluctuations of the ectodomain in the symmetric intermediate conformation with four insulins derived from GNM. The different domains of IR are labeled along the horizontal and vertical axes. F1, F2 and F3 are the abbreviations of Fnш-1, Fnш-2 (Fnш-2a and Fnш-2b) and Fnш-3 domains. F2^+^ refers to a complex structure containing Fnш-2a, ID-α, αCT, ID-β and Fnш-2b domains. Different regions in the figure are labeled for convenience of description. **D.** Disappearance order of the interface contacts derived from the iteratively application of GNM. The interface contacts disappear in thirteen batches, which takes place in these three regions in an alternate manner. The labels along horizontal axis are the residue index and the corresponding domains in one partner chain, and the ones along the vertical axis are the residue index and the corresponding domains in the other partner chain.

The normalized inter-residue cross-correlation fluctuations (NIRCCFs) are calculated to explore the relative movements of different individual domains within the ectodomain. As shown **Fig. 10C**, the positive NIRCCFs in the interior of L1∼CR/L1’∼CR’ domains (region ①/①’), L2/L2’ domains (region ②/②’), Fnш-1/ Fnш-1’ (F1/F1’) domains (region ③/③’), Fnш-2^+^∼Fnш-3/Fnш-2’^+^∼Fnш-3’ (F2^+^∼F3/F2’^+^∼F3’) domains (region ④/④’) indicate that rigidity dominates the movements of these domains. The L1-CR domains yield negative NIRCCFs with Fnш-2^+^-Fnш-3 domains (region ⑤) and L1’∼CR’ domains (region ⑥), and also yield weak negative NIRCCFs with L2’∼Fnш-1’-Fnш-2’^+^∼Fnш-3’ (L2’∼F1’∼F2’^+^∼F3’) domains (region ⑨). The Fnш-2^+^∼Fnш-3 (F2^+^∼F3) domains yield negative NIRCCFs with Fnш-2’^+^∼Fnш-3’ (F2’^+^∼F3’) domains (region ⑩). The negative NIRCCFs imply that the directions of the movements are opposite. The movements between the domains involved in region ⑨ and ⑩ facilitate the formation the complete T1BS (region ⑨) and the dimerization of Fnш-3-Fnш-3’ domains (region ⑩) which can further help the dimerization of the two tyrosine kinase domains. The positive NIRCCFs between L2∼Fnш-1 (L2∼F1) domains and L2’∼Fnш-1’ (L2’∼F1’) domains (region ⑧) are consistent with the structural features that interface contacts are present in the L2∼Fnш-1∼L2’∼Fnш-1’ dimer. The weak negative and positive correlations between L1’-CR’ domains and L2∼Fnш-1∼Fnш-2^+^∼Fnш-3 (L2∼F1∼F2^+^∼F3) domains (region ⑦) indicate the complexity of insulin-mediated interactions between the domains. Because insulin 1, 2, 1’ and 2’ (I1, I2, I1’ and I2’) make contacts with the L1 domain, Fnш-1 (F1) domain, L1’ domain and Fnш-1’ (F1’) domain, respectively, the NIRCCFs between the insulins and their interacting domains are positive.

The disappearance order of the interface contacts is calculated according to the distance fluctuations derived from the iterative application of GNM. As shown in **Fig. 10D**, the interface contacts disappear in thirteen batches. The first batch is composed of only one interface contact in the first sub-region of region 1, which locates at the left bottom of the interface (**Fig. 10B**). The second and third batch contain all the interface contacts in the second and third regions, respectively. All the other interface contacts are found in the four sub-regions of region 1, which disappear in the four sub-regions in an alternate manner. During the disappearance of the interface contacts (**Fig. S7A, S7B** and **S7C**), the positive NIRCCFs in the interior of the L2/L2’ domains increase gradually, whereas the positive NIRCCFs between L2∼Fnш-1 (L2∼F1) domains and L2’∼Fnш-1’ (L2’∼F1’) domains decrease gradually, and become weak negative at the end (after disappearance of all interface contacts between them). Because of the disruption of all interface contacts, the weak positive NIRCCFs between L2∼Fnш-1 (L2∼F1) domains and L2’∼Fnш-1’ (L2’∼F1’) domains become weak negative. The negative NIRCCFs between L1∼CR domains and Fnш-2^+^∼Fnш-3 (F2^+^∼F3) domains decrease, which may be related to the ensuing formation of the complete T1BS and dimerization of Fnш-3∼Fnш-3’ (F3∼F3’) domains.

As shown in **Fig. 11**, another T1BS in the asymmetric active state is not occupied by insulin. It appears that there is not enough space for insulin to be bound inside the pocket. But the T2BS on the Fnш-1 domain of the partner can bind an insulin in its asymmetric state (**Fig. 11**). This structural information suggests that the second complete T1BS may be formed in a different way. Binding of insulin to the structure around the T1BS in the asymmetric active state would induce structural changes in three parts (L1’ domain and the αCT and Fnш-1 domains of the partner) of the T1BS, resulting in the formation of the second compete T1BS. Because of its relatively-weak binding affinity (estimated by PRODIGY [33, 34]; results described later), insulin at the T2BS (shown in **Fig. 11**) may readily dissociate from this site on the Fnш-1 domain. As this insulin molecule is very close to the T1BS (shown in **Fig. 11**), it might interact directly with the L1’ domain and the αCT domain of the partner after dissociation, thereby inducing structural changes to form the complete T1BS. Certainly, it is also possible that another insulin molecule might come in and bind to the structure to induce similar conformational changes. When two complete T1BSs are formed and the IR’s overall structure is solidified with the binding of two insulin molecules to its two T1BSs, IR is in its fully-activated state along with the formation of tyrosine kinase dimer intracellularly, with the whole receptor structure being symmetric again.

**Figure 11.**
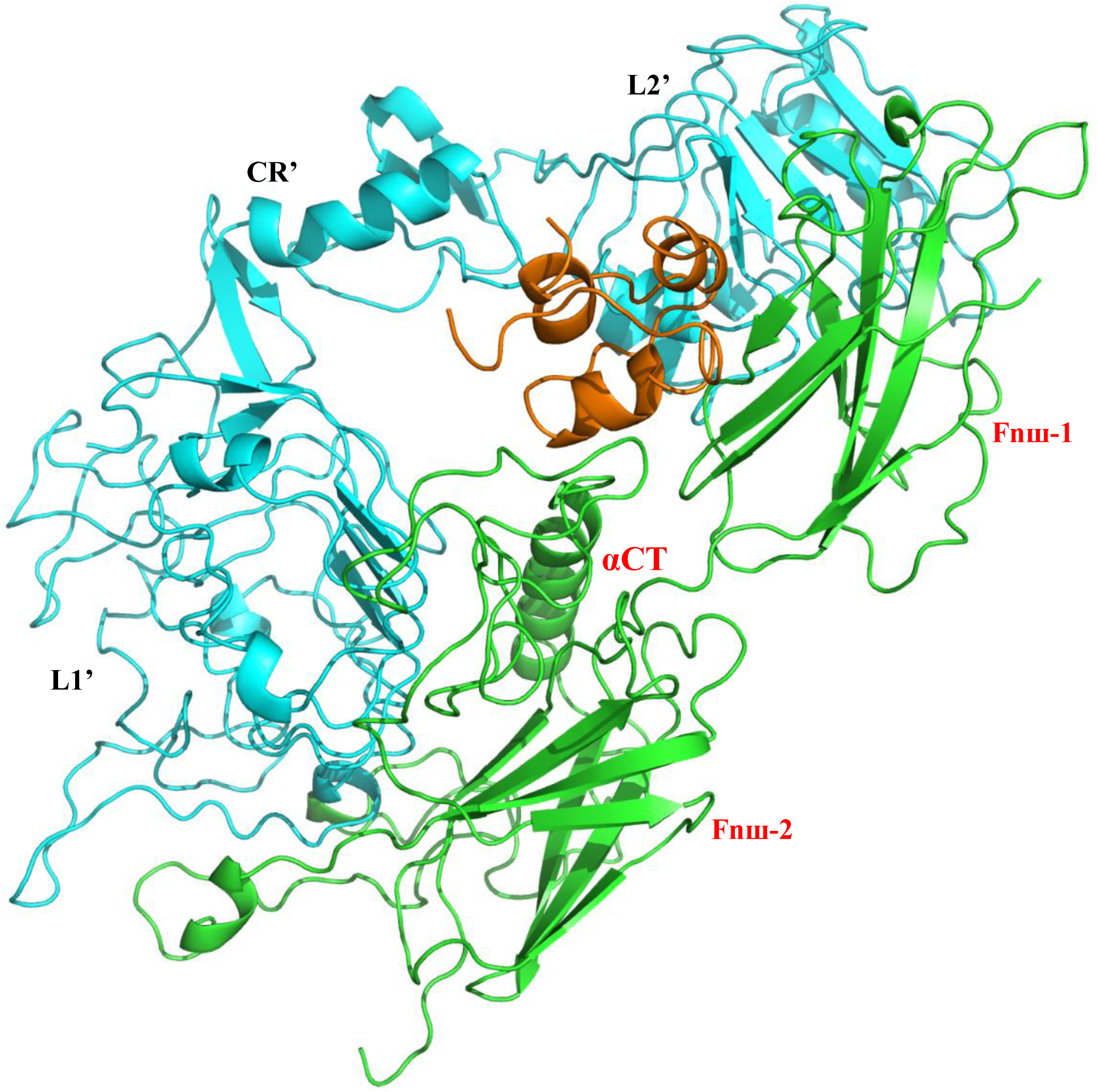
Incomplete T1BS in the representative asymmetric active conformation. The two chains of IRs are colored in green and cyan. Insulin at the T2BS is colored in orange.

Additionally, the structure of the homodimer formed by transmembrane (TM), juxtamembrane (JM), tyrosine kinase (TK) and βCT domains is shown in **Fig. S8A** (details of the construction process are described in earlier section). Based on the constructed conformational interaction between the βCT domain and the TK’ domain of the partner (**Fig. S8B**), it appears that the relative movements and interactions between the βCT domain (or the βCT’ domain) and the TK’ domain (or the TK domain) of the partner would facilitate the formation of the homodimer.

### Role and order of insulin binding to the two types of insulin binding sites

Based on the proposed structural models, the minimum and maximum numbers of insulins that exist in the representative conformations are summarized in **Table 2**. One or two insulin molecules may bind to the T2BS(s) in the auto-inhibited state first, which would result in structural rearrangement in the L1∼CR∼L2∼Fnш-1 domains. Then, another one or two insulin molecules would bind to the incomplete T1BS(s) in the intermediate state, which induces conformational changes and forms the complete T1BS(s). The conformational changes induced by the formation of one or two complete T1BS(s) are transmitted from the L1∼CR∼L2∼Fnш-1 and ID-α∼αCT∼ID-β domains to the Fnш-2/3, transmembrane and juxtamembrane domains, and this critical process might also facilitate the formation of the functional dimer of the tyrosine kinase domains. In the active states, the insulin molecules may remain in the T2BSs or may disassociate from these sites (**Fig. S3C, S3D, S3E, S3F**).

**Table 2.** Minimum and maximum numbers of insulins that may exist in the four representative IR conformations.

| State | Number of insulins at T1BS |  | Number of insulins at T2BS |  |
| --- | --- | --- | --- | --- |
|  | minimum | maximum | minimum | maximum |
| **Apo form (auto-inhibited state)** | 0 | 0 | 0 | 2 |
| **Symmetric intermediate state** | 1 (at incomplete T1BS) | 2 (at incomplete T1BS) | 0 | 2 |
| **Asymmetric active state** | 1 | 1 | 0 | 2 |
| **Symmetric active state** | 2 | 2 | 0 | 2 |

In this study, attempts are also made to predict the change of the relative binding affinities between insulin and the ectodomain of IR in the representative conformations by using PRODIGY, which is a contacts-based protein-protein binding affinity predictor [33, 34]. The results are summarized in **Table 3**. The binding affinity between insulin and its T2BS sequentially decreases from its apo form to intermediate state and then further to its active states. In contrast, the relative binding affinity between insulin and its T1BS increases during transition from its intermediate state (which contains the incomplete binding site) to the two representative active states (which contain the complete binding site). The earlier-reported “negative cooperativity” of insulin’s binding interaction with its T1BSs and T2BSs [35] might be understood on the basis of insulin-induced dynamic changes of the 3-D structures and conformations of the two types of insulin binding sites in the ectodomain.

**Table 3.** Relative binding affinity (*RBA*) between insulin and the ectodomain of IR in the four representative conformations predicted by PRODIGY [34].

| Conformation | Predicted *RBA* between insulin and the ectodomain of IR (Kcal/mol) |  |
| --- | --- | --- |
|  | T1BS | T2BS |
| Apo form (auto-inhibited state) | – | $-13.2 \pm 1.4$ |
| Symmetric intermediate state | $-8.7 \pm 0.8$ | $-11.5 \pm 0.3$ |
| Asymmetric active state | $-22.2$ | $-9.5 \pm 0.1$ |
| Symmetric active state | $-18.0 \pm 0.4$ | $-10.8 \pm 0.0$ |

## CONCLUDING REMARKS

In this work, it is proposed, on the basis of partial experimental structures available, that the full-length IR might exist in four representative conformations, which include: the apo form (auto-inhibited state), the symmetric intermediate state, the asymmetric active state, and the symmetric active state. The full structures of the complete ectodomain in these four states are constructed using computational modeling approach. According to the four representative conformations, the symmetric and asymmetric conformational changes are analyzed to deduce the transition pathway during IR activation. The proposed conformational transition pathway suggests that the conformational changes induced by insulins at the two types of binding sites play different roles during the process of IR activation. Insulin binding to T2BS(s) would induce structural rearrangements and lead to the exposure of the incomplete T1BS(s). Subsequent insulin binding to the incomplete T1BS(s) would further induce additional conformational changes to form the active state. If two complete T1BSs are formed in an sequential order, then an asymmetric conformational transition would take place. It is likely that one insulin at the complete T1BS might be sufficient to drive the receptor into the active state; however, the binding of two insulins at the two T1BSs would solidify the receptor conformation to form a more stable, symmetric active state. With the formation of the final symmetric active state, insulin molecules at the T2BSs likely would dissociate from the site(s) because of reduced binding affinities as a result of conformational changes. Lastly, dissociation of insulins from the two T1BSs would drive the IR conformation back to its original inactive state (*i.e.*, the apo form).

## MATERIALS AND METHODS

### Sequence and experimental structures of human IR

There are 1382 amino acids in human IR protein structure [5]. Based on the information presently available in the public domains, it appears that all experimentally-determined structures of IR are incomplete. For convenience, the basic information of the main experimental structures of human IR used in this work is listed in **Table S1**; their structures are shown in **Fig. S3**; and ranges of amino acid residues, full names and abbreviations for different domains are listed in **Table S2.** There are two types of insulin binding sites in the IR structure [17, 18]: type 1 binding site (T1BS) and type 2 binding site (T2BS). The complete T1BS is composed of L1 domain and the αCT’ and Fnш-1’ domains of the partner. T2BS is on side of Fnш-1 domain. In **Fig. S3A**, the incomplete T1BS of insulin on the L1/1’ domains are partially or completely hidden in the receptor dimer, but the T2BS on the Fnш-1 domain are exposed to the solvent [10–12]. The incomplete T1BS on the L1/1’ domains with two insulins bound inside are shown in **Fig.S3B** [13, 14], and the complete T1BS structures are shown in **Fig. S3C, S3D, S3E** and **S3F** [15–18]. The two T2BSs with insulin molecules bound inside are shown in **Fig. S3F** [17, 18]. The transmembrane and tyrosine kinase domains are shown in the left and right panel of **Fig. S3G**, respectively [19, 20]. Additionally, the structure of the complete insulin molecule is shown in **Fig. S3H** [28].

Based on the incomplete experimental structures of different parts of IR, a complete structure is assembled. Different domains of IR are shown in **Fig. S2**. While **Fig. S2A** shows the full-length IR (one chain), **Fig. S2B‒S2O** shows the signal peptide, L1 (leucine-rich repeat domain 1), CR (cysteine-rich region), L2 (leucine-rich repeat domain 2), Fnш-1 (fibronectin type-ш domain 1), Fnш-2a (fibronectin type-ш domain 2a), ID-α (insert domain of the α chain), αCT (C-terminal domain of the α chain), ID-β (insert domain of the β chain), Fnш-2b (fibronectin type-ш domain 2b), Fnш-3 (fibronectin type-ш domain 3), TM and JM (transmembrane domain and juxtamembrane domain, respectively), TK (tyrosine kinase), and βCT (C-terminal domain of the β chain).

### Computational modeling

In this study, the experimental structures are assembled into the full-length IR in their corresponding states. The missing residues in all the structures are predicted and added using the SWISS-MODEL (https://swissmodel.expasy.org/), which is a web-based tool for homology modelling [23]. If there are no templates available for the missing sequence segments, contact-based protein structure prediction is adopted to generate the corresponding structural segments. The SPOT-1D [25] and SPOT-Contact methods [26] are used to predict the protein secondary structure and residue-residue contact map, respectively. The Confold2 [24], a contact-guided protein structure prediction method, is employed to construct the protein three-dimensional (3-D) structures. The missing local segments or domains in one state are supplemented according to their corresponding structures and relative positions which exist in another state. VMD (Visual Molecular Dynamics) [22] is used to superimpose the common local structures and place the missing local segments or domains to the proper position.

The possible intermediate states along the conformation transition pathways are generated using structural alignment and rotation with VMD [22]. All the structures in different states are optimized to avoid clashes between atoms. The topology files are generated using CHARMM-GUI (http://www.charmm-gui.org) [29], and the CHARMM36m force field [36] is adopted. Energy minimization in vacuum is conducted using NAMD [30].

### Gaussian network model

The relative movements between different domains are investigated using Gaussian network model (GNM) which is a simplified physical model for protein dynamics [31]. In GNM, each amino acid in protein is represented by the Cα atom. Two Cα atoms would be connected with a spring when the distance between them in the experimental structure is equal to or smaller than a given cutoff (in this study 7 Å is adopted, which includes all pairs of residues within a first inter-residue coordination shell [32]). The internal Hamiltonian of the protein system can be written as [37]

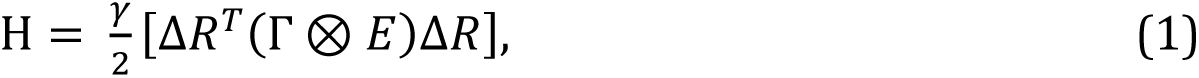

where γ is the spring contants, Δ*R* is the 3*N*-dimensional displacement vectors for the fluctuations of the Cα atoms, and *N* is the number of the residues in the protein system. ΔR*^T^* is the transpose of Δ*R*, Δ*R^T^* = {Δ*X1*, Δ*Y1*, Δ*Z1*, Δ*X2*, Δ*Y2*, Δ*Z2*, …, Δ*XN*, Δ*YN*, Δ*ZN*}. *E* is the identity matrix with order 3, and ⊗ is the direct product. Γ is the Kirchhoff (or connectivity) matrix. Γ*ij* is defined as [37]

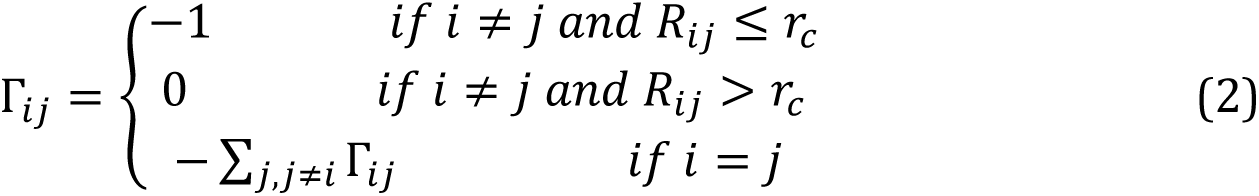

where *Rij* is the distance between the *i*^th^ and *j*^th^ Cα atoms in experimental structure, and *r_c_* is the cutoff.

The mean square fluctuations of the *i*^th^ residue and the mean cross-correlation fluctuations between the *i*^th^ and *j*^th^ residues can be calculated as [31]

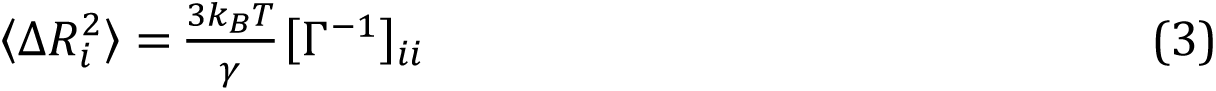

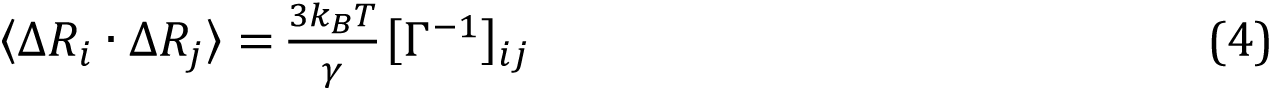

where Δ*Ri* is the displacement vector {Δ*Xi*, Δ*Y*i, Δ*Z*i} of the *i*^th^ residue, *kB* is Boltzmann constant, T is the absolute temperature, and γ is the spring constant. Γ^‒1^ is the pseudo-inverse of the Γ matrix.

The cross-correlation fluctuations is normalized as [32]

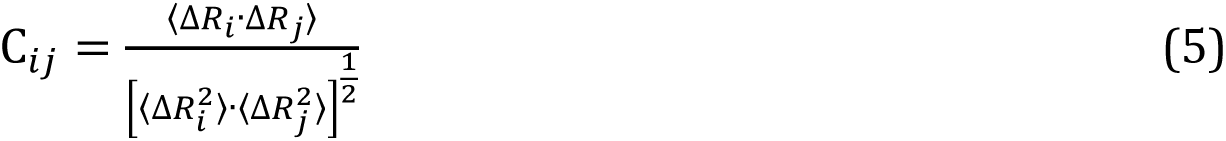

The relationship between experimental Debye-Waller or B-factor and the mean square fluctuation of the *i*^th^ residue can be expressed as [32]

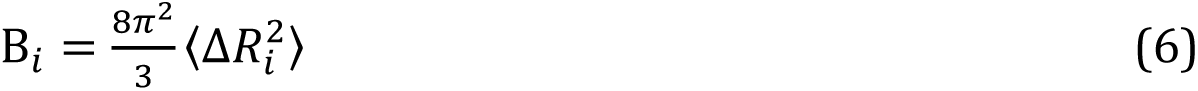

The mean square fluctuation of the distance vector between the *i*^th^ and *j*^th^ residues can be calculated as [38]

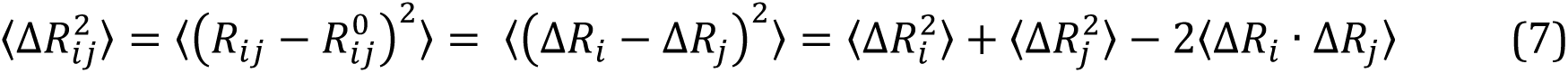

In the work conducted by Su *et al.* [39], protein unfolding behavior is studied by iterative application of GNM. The native contact with the largest distance fluctuation would be broken first as the temperature is gradually increased [39]. Here, the disassociation of the two partners in the ectdomain of IR is simulated in a similar way for protein unfolding proposed by Su *et al.* [39]. The native interface contacts between the two partners disappear in order according to the magnitudes of the mean square fluctuations in the distance vectors. If the strength of the interface interaction between the *i*^th^ and *j*^th^ residues is weaker, the interface contact vanishes more easily. The mea square fluctuation in the distance vector between *i*^th^ and *j*^th^ residues can reflect the strength of the interaction.

### Binding energy calculations

The binding energies/affinities between insulins and the ectodomains of the full-length IR are evaluated using PRODIGY ((PROtein binDIng enerGY prediction), a contact-based protein-protein binding affinity predictor, which is developed by Vangone and Bonvin [33, 34]. Number of interface contacts and non-interacting surface areas are adopted in the linear function for calculating the protein-protein binding affinity based on the structure of protein-protein complex [33]. PRODIGY is downloaded from the website (https://github.com/haddocking/prodigy) and set up on local machine [34].

## LEGENDS FOR SUPPLEMENTARY FIGURES

**Supplementary Figure S1.**
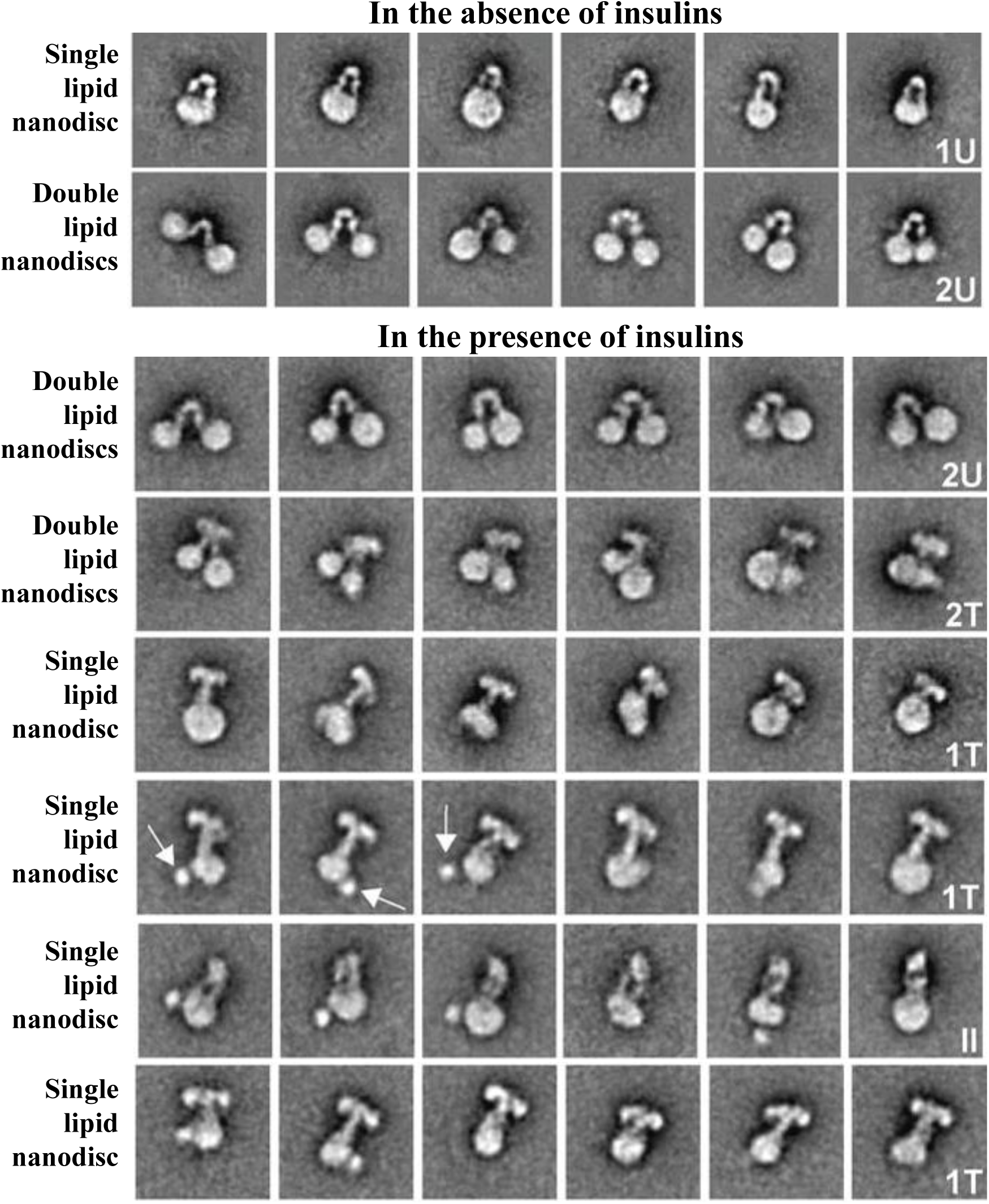
Different groups of cryo-EM images of the full-length IR obtained by Gutmann *et al.* [21]. In the absence of insulins, the full-length IR presents as (inverted) U-shaped conformations in single or double lipid nanodiscs. In the presence of insulins, the full-length IR presents as U-shaped or T-shaped conformations in double lipid nanodiscs, and T-shaped or II-shaped conformations in single lipid nanodisc.

**Supplementary Figure S2.**
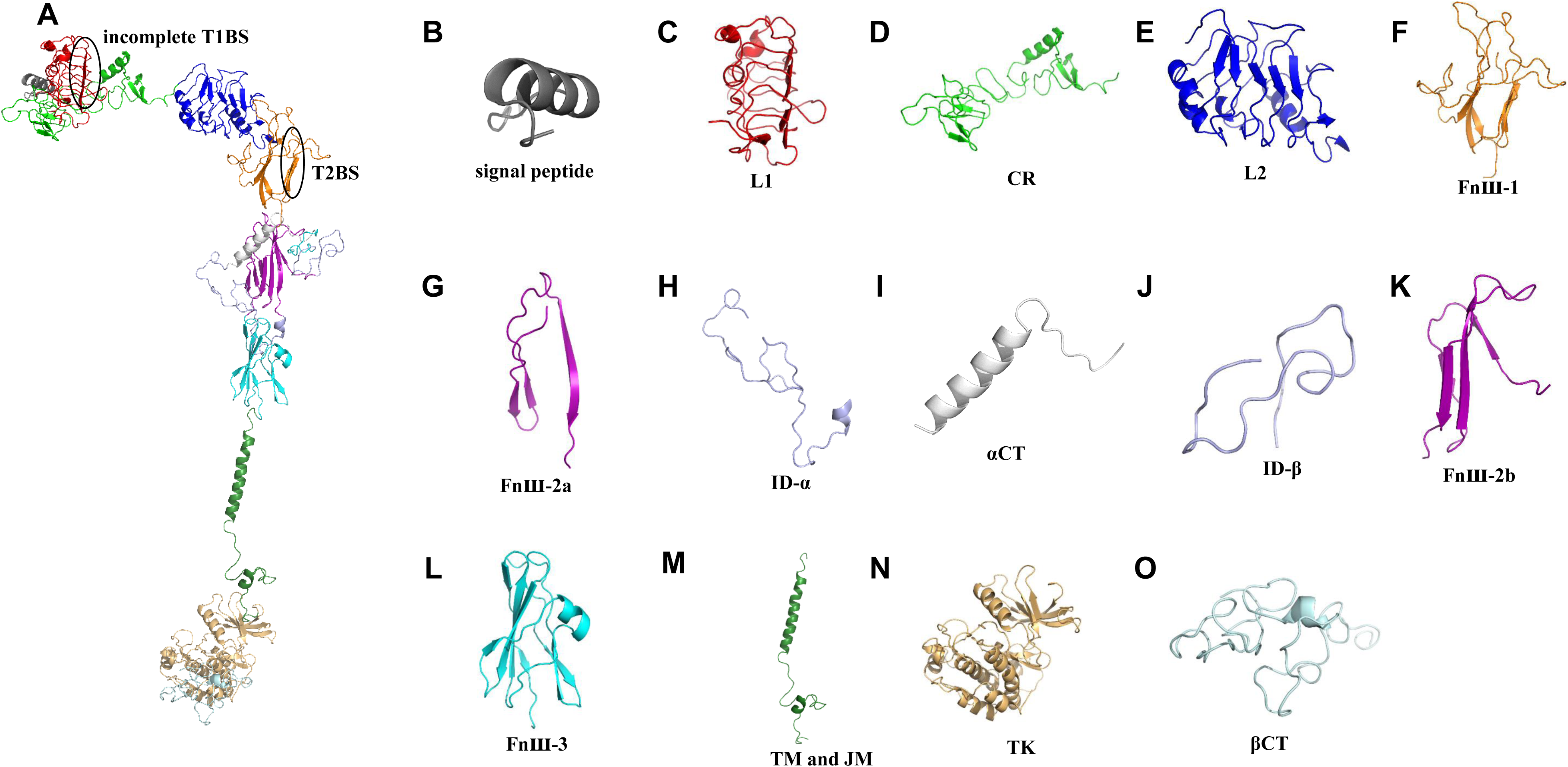
Different domains of the full-length IR. **A.** Structure o full-length IR (one chain, 1382 amino acids). **B.** Signal peptide (residues 1‒27, gray color). **C.** L1 (leucine-rich repeat domain 1, residues 28‒183, red color)**. D.** CR (cysteine-rich region, residues 184‒337, green color). **E.** L2 (leucine-rich repeat domain 2, residues 338‒497, blue color). **F.** Fnш-1 (fibronectin type-ш domain 1, residues 498‒622, orange color). **G.** Fnш-2a (fibronectin type-ш domain 2a, residues 623‒665, purple color). **H.** ID-α (insert domain of the α chain (residues 1-750), residues 666‒719, light blue color). **I.** αCT (C-terminal domain of the α chain, residues 720‒746, white color). **J.** ID-β (insert domain of the β chain (residues 751‒1382), residues 763‒792, light blue color). **K.** Fnш-2b (fibronectin type-ш domain 2b, residues 793‒847, purple color). **L.** Fnш-3 (fibronectin type-ш domain 3, residues 848‒947, cyan color). **M.** TM and JM (transmembrane and juxtamembrane domains, residues 948‒1018, forest color). **N.** TK (tyrosine kinase, residues 1019-1295, light orange color). **O.** βCT (C-terminal domain of the β chain, residues 1296‒1382, palecyan color).

**Supplementary Figure S3.**
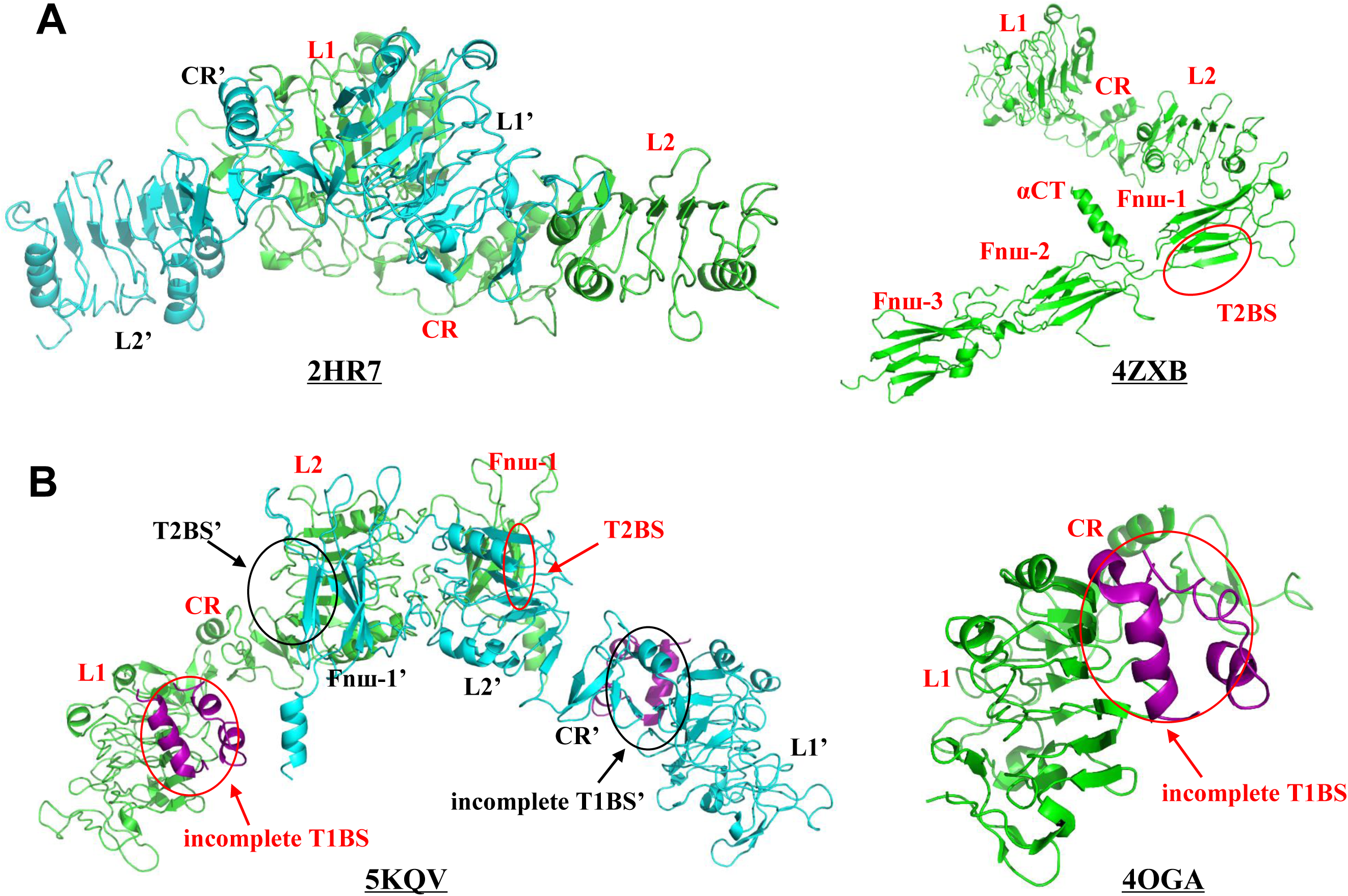
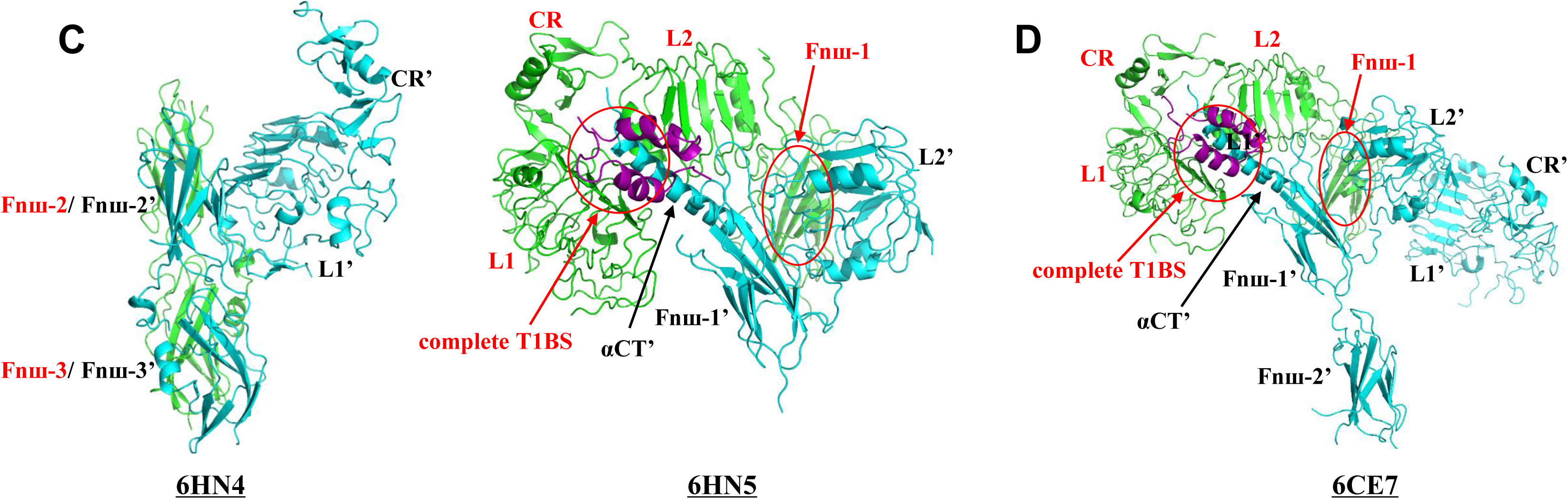
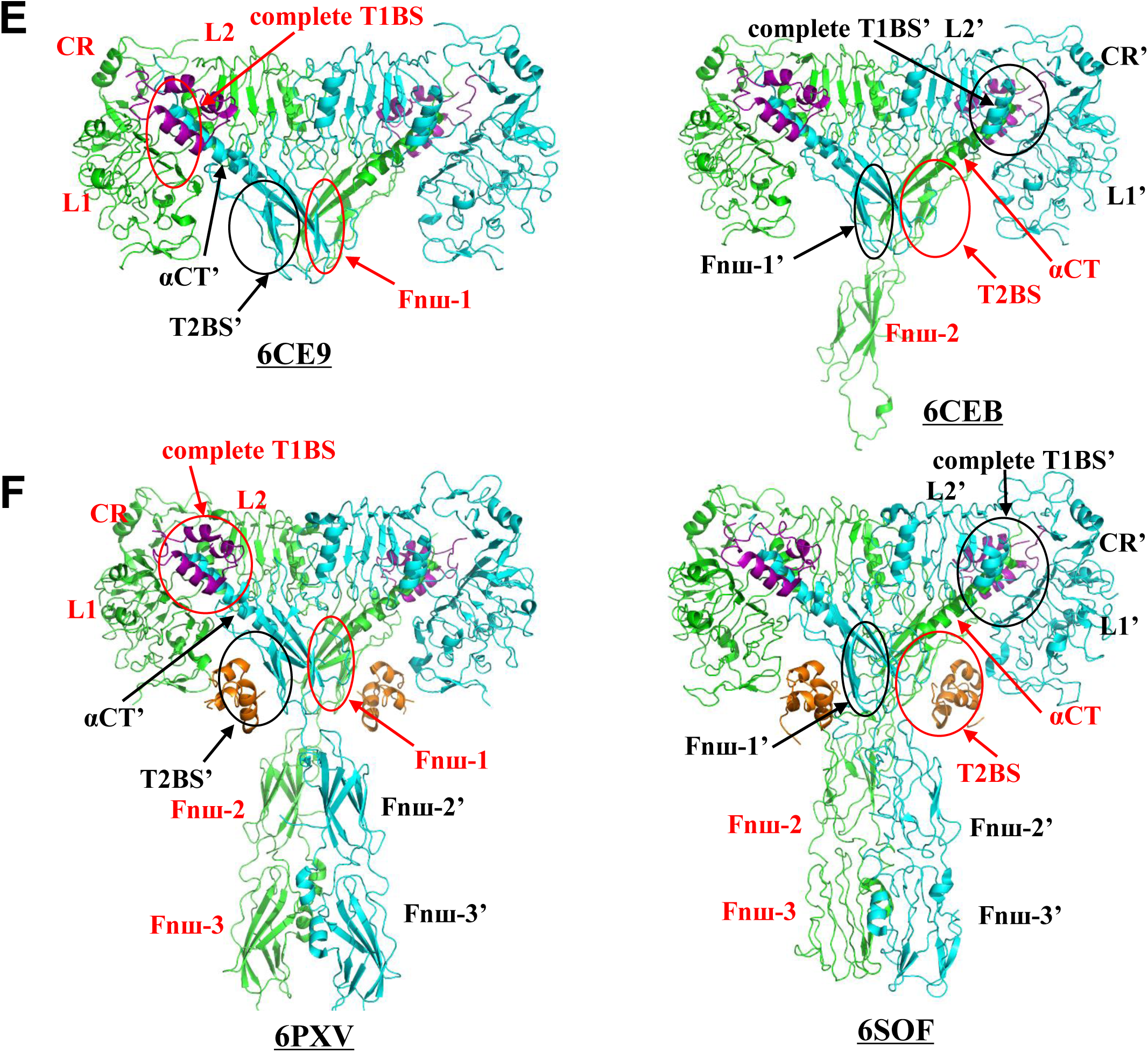
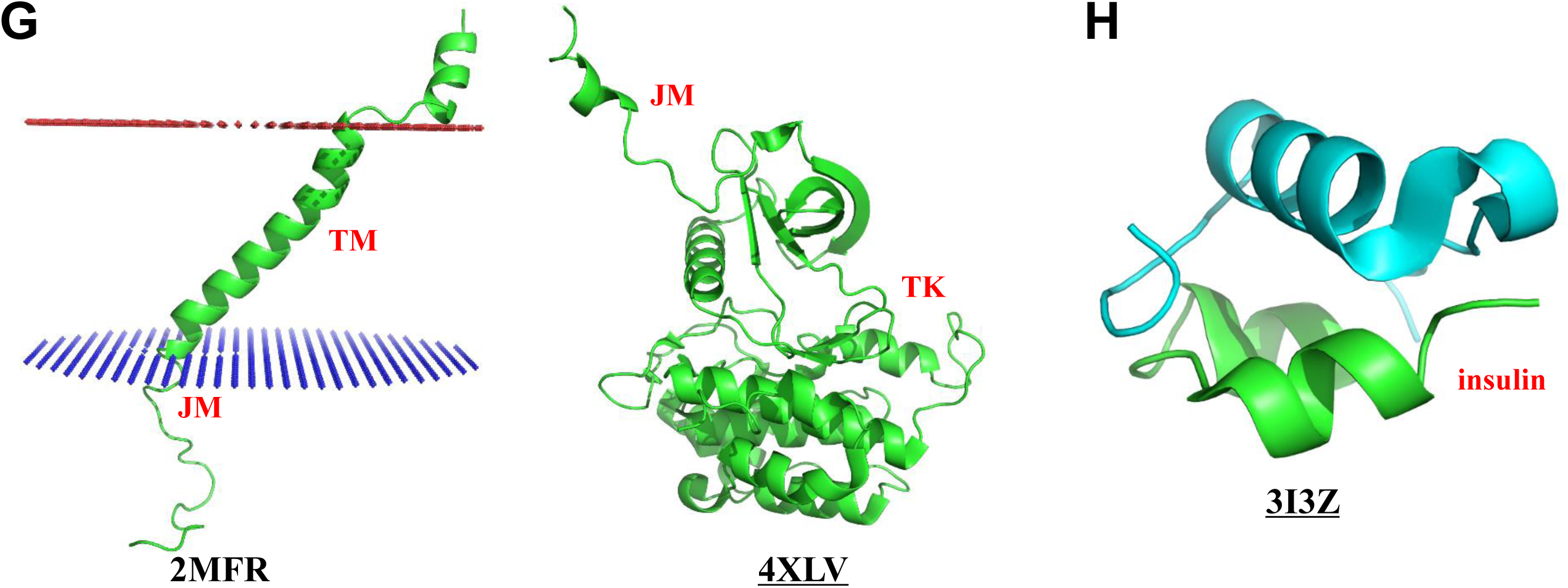
Experimental structures of human insulin receptor (IR) and insulin. **A-F**. The ectodomain of IR. **A.** The apo form or auto-inhibited state (PBD code: 2HR7, 4ZXB). **B.** Symmetric intermediate state with two insulins (PBD code: 5KQV, 4OGA). **C.** Asymmetric state 1 with one insulin (PDB code: 6HN4, 6HN5). **D.** Asymmetric state 2 with one insulin (PDB code: 6CE7). **E.** Symmetric state with two insulins (PDB code: 6CE9, 6CEB). **F.** Symmetric state with four insulins (PDB code: 6PXV, 6SOF). **G.** Transmembrane domain (left panel; 2MFR) and tyrosine kinase domain (right panel; 4XLV). **H.** Insulin (3L3Z). Note that all basic information of these structures is summarized in **Table S1**, and the full names and abbreviations of different domains are listed in **Table S2.** Each pair of arrow and ellipse in red points to a domain in one partner chain which is colored in green; likewise, the pair of arrow and ellipse in black points to a domain in the other partner chain which is colored in cyan.

**Supplementary Figure S4.**
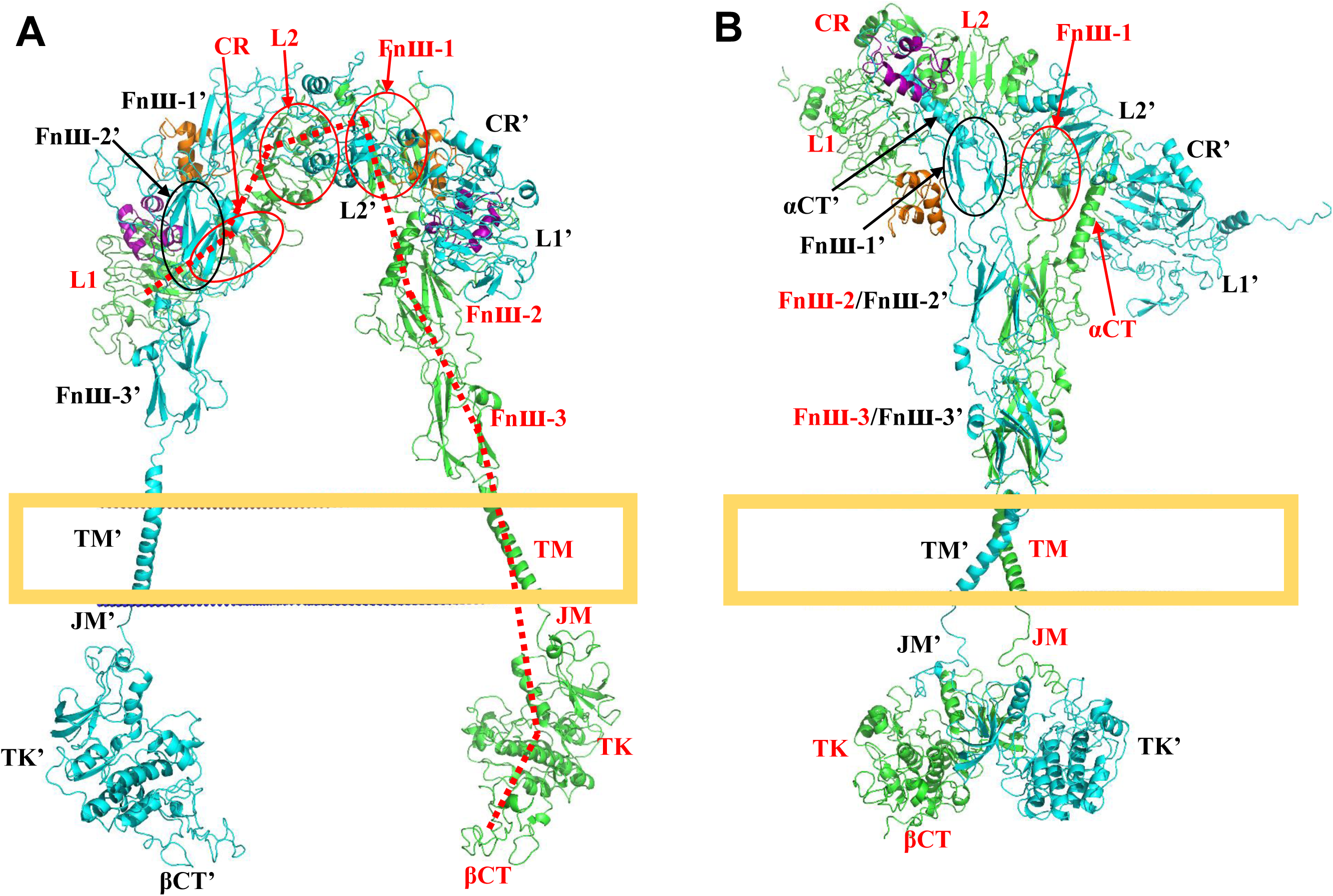
Conformations of the full-length IR at the symmetric intermediate state and a different asymmetric active state. **A.** Different view of the conformation at the symmetric intermediate state (also a U-shaped conformation). **B.** A asymmetric active state (PDB code: 6CE7, 6HN4, 6HN5, 4ZXB, 2MFR, 4XLV) that is slightly different from the asymmetric active state shown in **Figure 2C.** The insulins at the incomplete or complete T1BSs are colored in purple, and insulins at the T2BSs colored in orange. Each pair of arrow and ellipse in red points to a domain in one partner chain which is colored in green; likewise, the pair of arrow and ellipse in black points to a domain in the other partner chain which is colored in cyan.

**Supplementary Figure S5.**
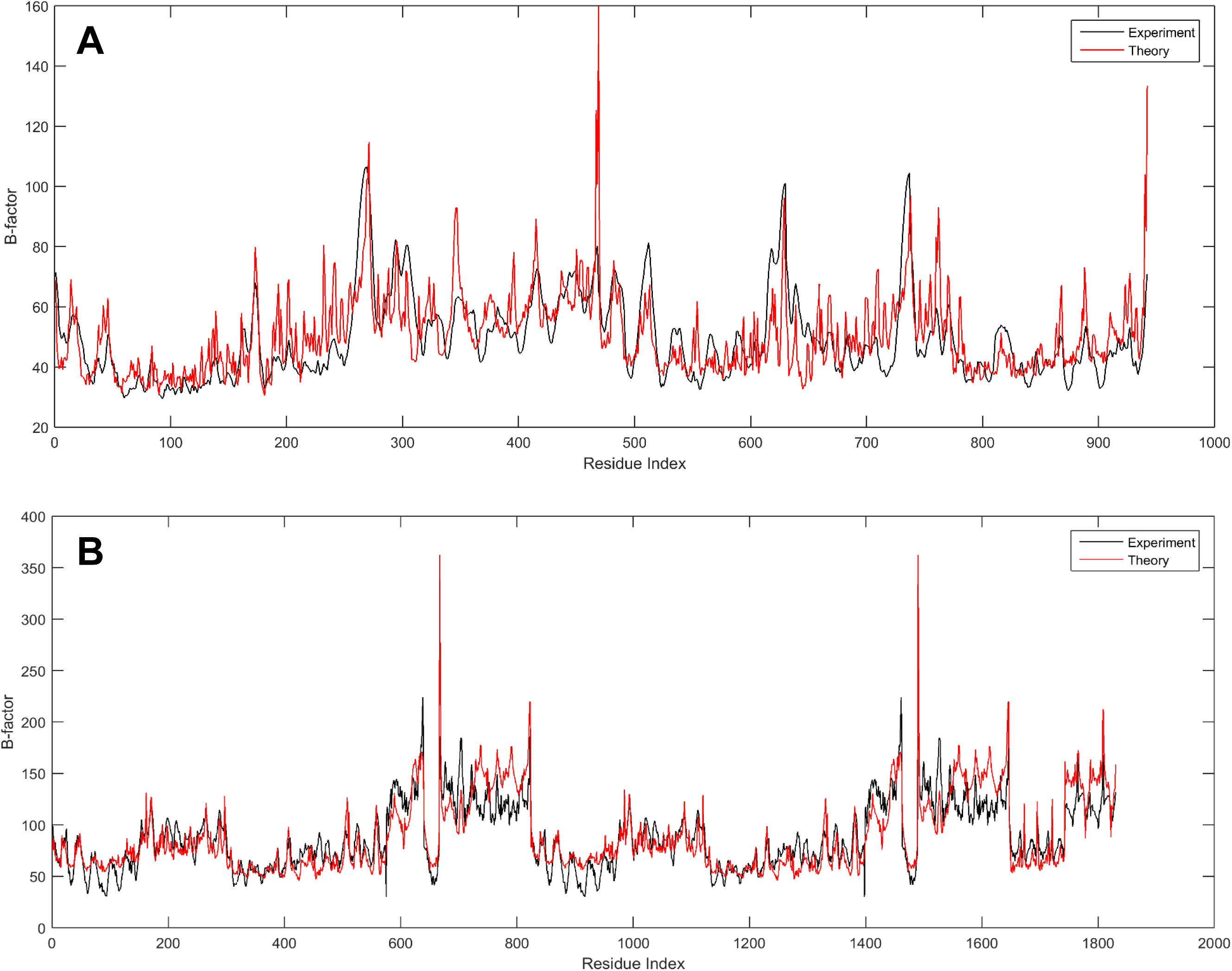
The experimental and theoretical B-factors of the L1∼CR∼L2∼L1’∼CR’∼L2’ dimer at the auto-inhibited state (PDB code: 2HR7) and the ectodomain at active state (PDB code: 6PXV). **A.** Experimental and theoretical B-factors of the L1∼CR∼L2 dimer at the auto-inhibited state (PDB code: 2HR7). The correlation coefficient between the experimental and theoretical B-factors of the Cα atoms is approximately 0.67. **B.** Comparison of the experimental and theoretical B-factors of the ectodomain at its active state (PDB code: 6PXV). The correlation coefficient is approximately 0.83 between the B-factors of the Cα atoms in experimentally-determined structure (in black lines) and the corresponding theoretical B-factors derived from GNM (in red lines).

**Supplementary Figure S6.**
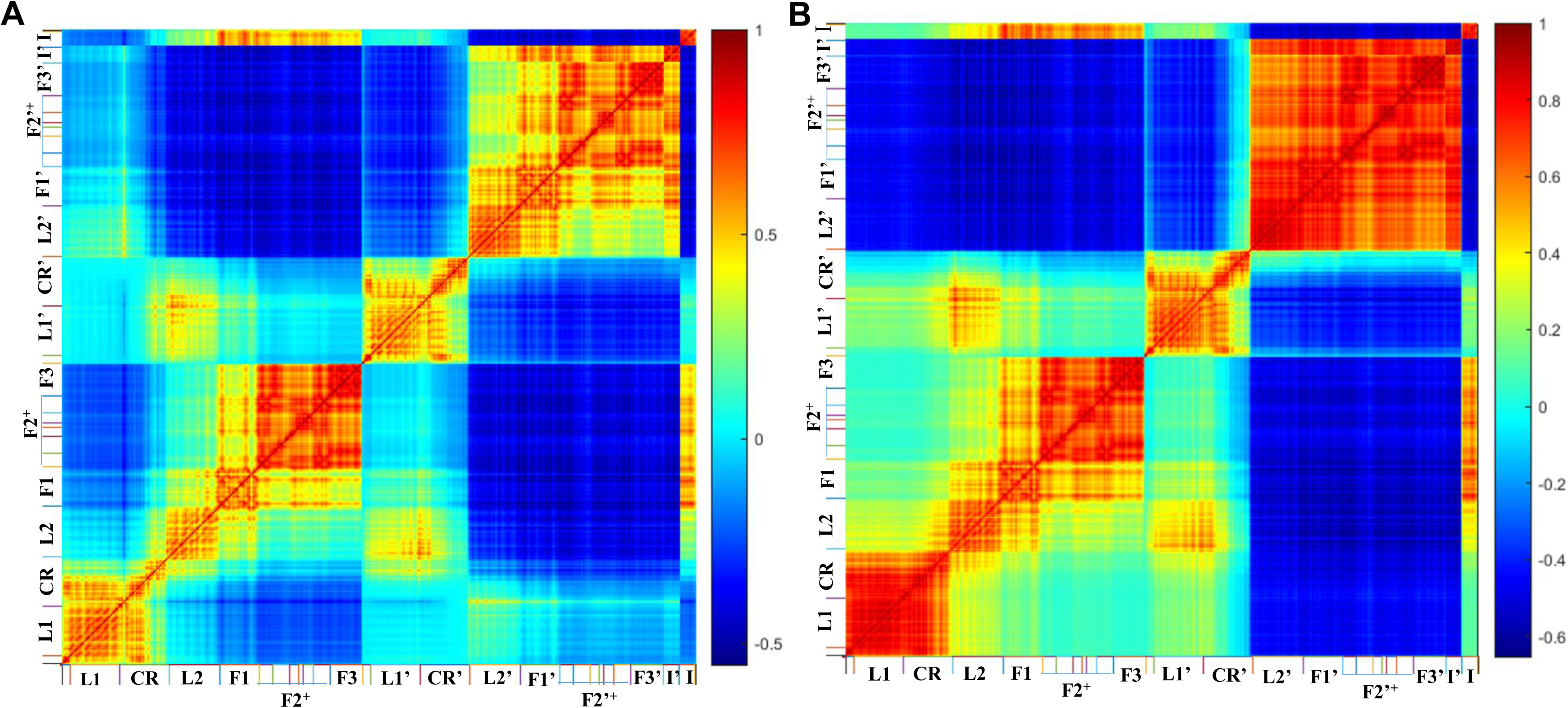
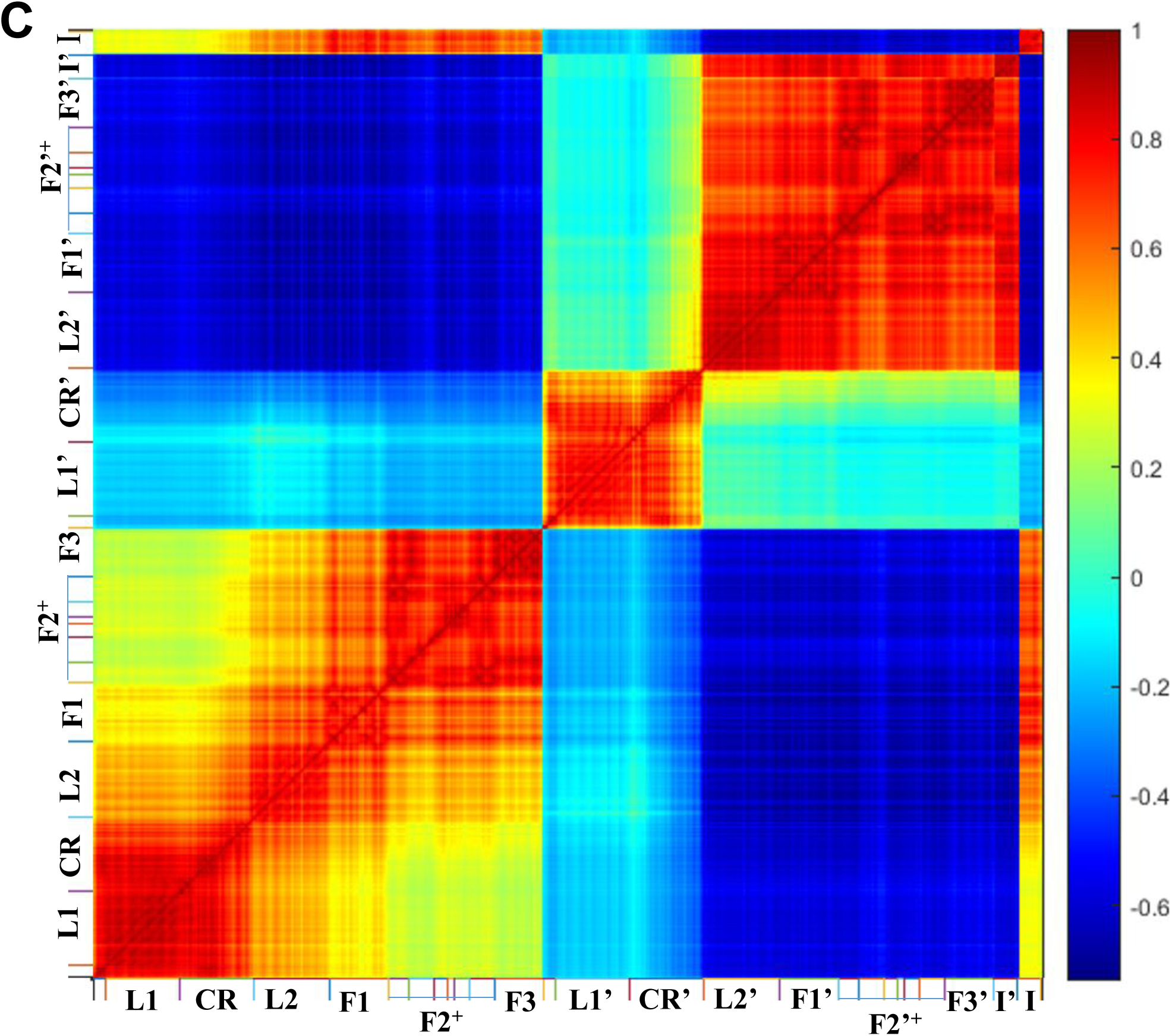
Normalized inter-residue cross-correlation fluctuations of the ectodomain in the auto-inhibited conformation with two insulins at the T2BSs during the vanishing process of interface contacts. **A.** Intermediate state after the disappearance of the interface contacts in the first and second regions. The loss-number-of-interface-contact (LNIC) is 9. **B.** Intermediate state after the disappearance of the interface contacts in the first, second and third regions (LNIC is 29). C. Intermediate state after the disappearance of the most interface contacts (LNIC of 51). F1, F2 and F3 are the abbreviations of Fnш-1, Fnш-2 (Fnш-2a and Fnш-2b) and Fnш-3 domains, respectively. F2+ refers to a complex structure containing Fnш-2a, ID-α, αCT, ID-β and Fnш-2b domains.

**Supplementary Figure S7.**
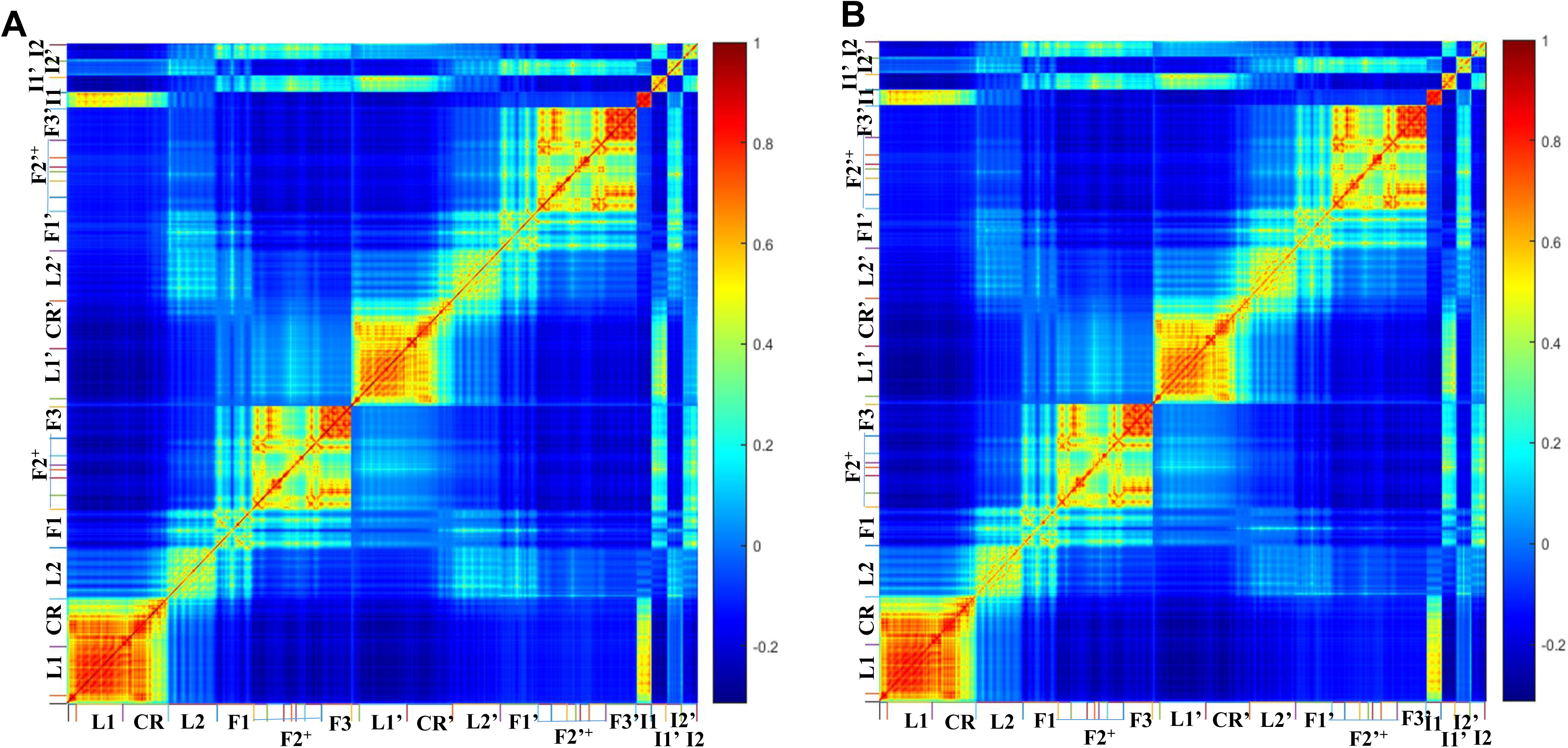
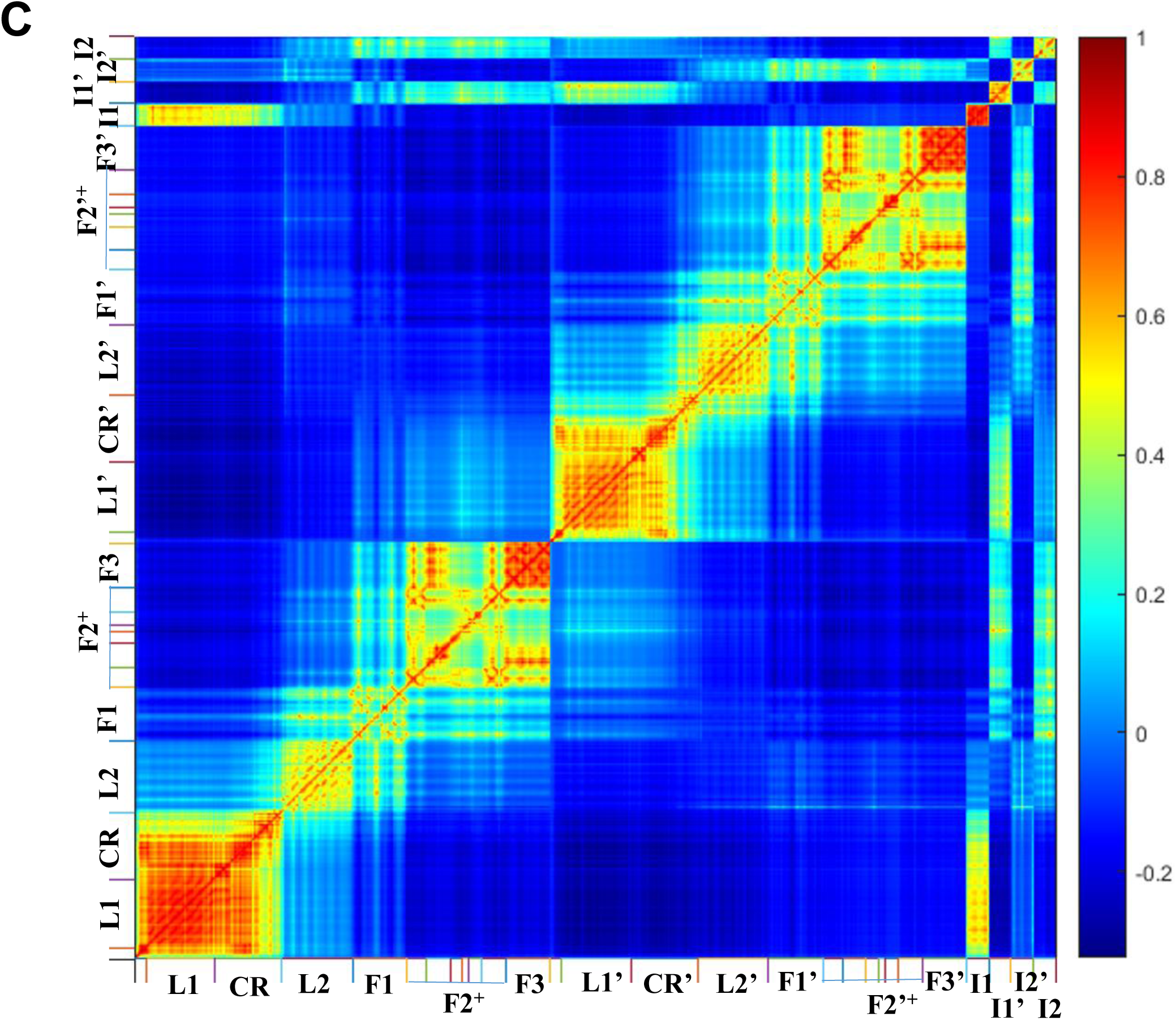
Normalized inter-residue cross-correlation fluctuations of the ectodomain in the symmetric intermediate conformation with four insulins during the vanishing process of interface contacts. **A.** Intermediate state after the disappearance of the interface contacts in the second and third regions. The loss-number-of-interface-contact (LNIC) is 6. **B.** Intermediate state after the disappearance of the interface contacts in the second, third and the firs 1.1 sub-regions (LNIC of 20). C. The state after disappearance of all interface contacts (LNIC of 25). F1, F2 and F3 are the abbreviations of Fnш-1, Fnш-2 (Fnш-2a and Fnш-2b) and Fnш-3 domains, respectively. F2+ refers to a complex structure containing Fnш-2a, ID-α, αCT, ID-β and Fnш-2b domains.

**Supplementary Figure S8.**
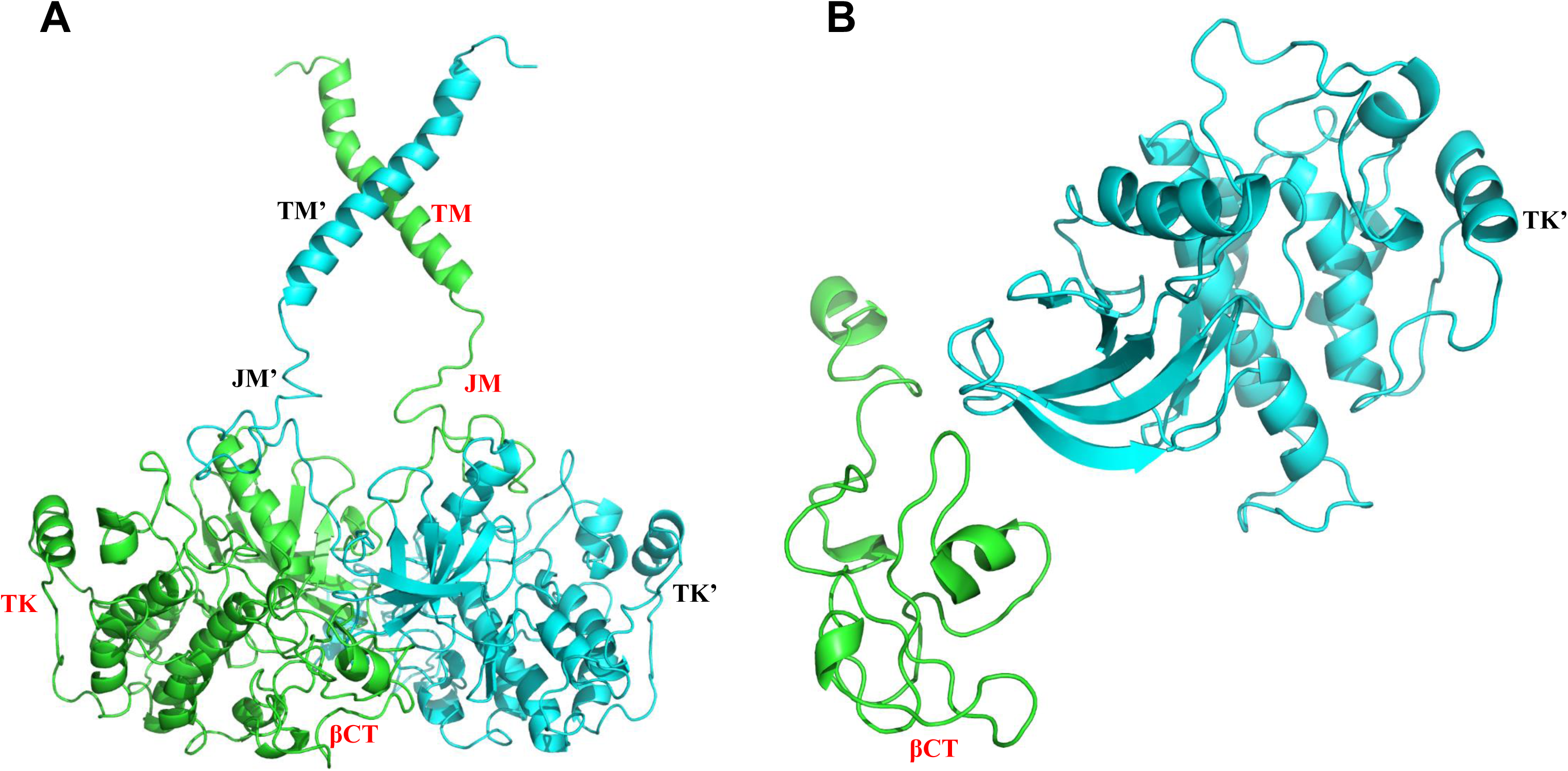
Structures of the functional dimer of the transmembrane (TM), juxtamembrane (JM), tyrosine kinase (TK) and βCT domains. **A.** Complete structure of the active dimer of TM, JM, TK and βCT domains. **B.** Possible conformations between TK domains and the βCT domain of the partner. The two chains of IRs are colored in green and cyan.

**Supplementary Table S1.** Experimental structures of IR used in this study.

| **Domain of IR** | **PDB code** | **Methods** | **Resolution (Å)** | **Number of insulins** | **Published Year** | **References** |
| --- | --- | --- | --- | --- | --- | --- |
| **Ectodomain** | 2HR7 | X-ray crystallography | 2.32 | 0 | 2006 | [10] |
|  | 4ZXB |  | 3.30 | 0 | 2016 | [11, 12] |
|  | 5KQV |  | 4.40 | 2 | 2013 | [13] |
|  | 4OGA |  | 3.50 | 1 | 2014 | [14] |
|  | 6HN4 | Cryo-EM | 4.20 | 0 | 2018 | [15] |
|  | 6HN5 |  | 3.20 | 1 | 2018 | [15] |
|  | 6CE7 |  | 7.40 | 1 | 2018 | [16] |
|  | 6CE9 |  | 4.30 | 2 | 2018 | [16] |
|  | 6CEB |  | 4.70 | 2 | 2018 | [16] |
|  | 6PXV |  | 3.20 | 4 | 2019 | [18] |
|  | 6SOF |  | 4.30 | 4 | 2020 | [17] |
| **Transmembrane domain (TD)** | 2MFR | NMR | – | – | 2014 | [19] |
| **Tyrosine kinase (TK)** | 4XLV | X-ray crystallography | 2.30 | – | 2015 | [20] |
| **Insulin** | 3I3Z | X-ray crystallography | 1.60 | – | 2010 | [28] |

**Supplementary Table S2.** Different domains of the full-length IR.

| Name of domain |  | Amino acid sequence |
| --- | --- | --- |
| Abbreviation | Full name |  |
| **SG** | Signal peptide | 1–27 |
| **L1** | Leucine-rich repeat domain 1 | 28–183 |
| **CR** | Cysteine-rich region | 184–337 |
| **L2** | Leucine-rich repeat domain 2 | 338–497 |
| **F$\text{III}$-1** | Fibronectin type-$\text{III}$ domain 1 | 498–622 |
| **F$\text{III}$-2a** | Fibronectin type-$\text{III}$ domain 2a | 623–665 |
| **ID-α** | Insert Domain of the $\alpha$ chain (residue 1-750) | 666–719 |
| **αCT** | C-Terminal domain of the $\alpha$ chain | 720–746 |
| **ID-β** | Insert Domain of the $\beta$ chain (residue 751~1382) | 763–792 |
| **F$\text{III}$-2b** | Fibronectin type-$\text{III}$ domain 2b | 793–847 |
| **F$\text{III}$-3** | Fibronectin type-$\text{III}$ domain 3 | 848–947 |
| **TM and JM** | Transmembrane and juxtamembrane domains | 948–1018 |
| **TK** | Tyrosine kinase domain | 1019–1295 |
| **βCT** | C-terminal domain of the $\beta$ chain | 1296–1382 |

